# Operant House: A Versatile Open-Source Platform for Automated Operant Conditioning Testing of Mice in Home Cages

**DOI:** 10.1101/2025.01.24.634815

**Authors:** Shintaro Otsuka, Yuriko Nakamura, Aya Ito-Ishida, Kunimichi Suzuki, Ayako Ishikawa, Keiko Matsuda, Shigetomo Suyama, Anis Contractor, Michisuke Yuzaki

**Author notes:** These authors contributed equally to this work (Shintaro Otsuka, Yuriko Nakamura, Aya Ito-Ishida).

## Abstract

Operant conditioning is a valuable method for studying cognitive functions, yet its adoption is limited by low throughput, labor intensity, and high costs. Here, we developed “Operant House,” a low-cost, programmable device featuring a touchscreen, retractable levers, and a water reward port, designed for flexible, automated operant conditioning tasks. To validate its utility, we implemented two protocols to assess working memory in mice: a delayed non-match-to-position test and a two-choice spatial discrimination test. Using these protocols, we examined a mouse model of Alzheimer’s disease carrying familial Alzheimer’s disease-associated amyloid precursor protein mutations. Results revealed significant working memory deficits as early as 5 months of age. These findings highlight the Operant House as a cost-effective, high-throughput platform for evaluating higher cognitive functions in mice, offering an accessible tool for investigating models of neurological and neuropsychiatric disorders.

## Introduction

Operant conditioning has long been utilized in behavioral neuroscience to evaluate cognitive performance in a variety of species. In these tests, subjects are trained to perform specific behaviors, such as touching panels or pressing levers, in order to obtain rewards, such as water and food. Well-established protocols have been developed to measure various aspects of cognitive functions, including visual and spatial recognition, attention, impulsivity, working memory, and flexibility (Dudchenko, 2004; Yu et al., 2009; Horner et al., 2013; Aoki et al., 2015; Kwak et al., 2015). Operant conditioning serves as an unbiased assay, providing valuable insights into mouse cognition within the context of various neuropsychiatric and neurological disease models, including schizophrenia, autism spectrum disorders, and Alzheimer’s disease. Conventional operant conditioning is typically conducted individually in a specialized apparatus under the supervision of an experimenter (Rossi and Yin, 2012). This process is labor-intensive and inefficient, as it must be repeated for each animal in a large cohort. Additionally, removing animals from their home cages—often during their inactive daytime period—for daily training sessions can induce significant stress. These limitations present substantial challenges to the widespread application of operant conditioning in disease model animals.

To overcome these challenges, various devices have been developed to enable automated operant conditioning within the mouse’s home cages, particularly during the correct light cycle for the mouse’s wakeful period. For example, a home cage that connects to a commercial operant chamber device for the 5-choice serial reaction time task were shown to achieve a tenfold increase in learning speed compared to conventional chambers for operant conditioning (Remmelink et al., 2017). However, conducting simultaneous tests on multiple mice using such equipment is challenging due to its high cost and large size. An alternative approach is the IntelliCage system, which allows a group of individually tagged mice to be housed together in a single large cage for operant conditioning (Codita et al., 2010; Endo et al., 2011; Voikar et al., 2018; van Dijk et al., 2019; Kiryk et al., 2020; Iman et al., 2021). While the IntelliCage provides valuable population-level behavioral information, individual mouse behavior can be influenced by the presence of other mice in the same cage (Benner et al., 2014; Benner et al., 2015; Ujita et al., 2018; Horigane et al., 2020). Therefore, there is a need for a more cost-effective and efficient method to perform operant conditioning in individual mice.

To address these issues, we developed a compact and affordable operant conditioning device, referred to as the “Operant House.” The Operant House is a fully programmable device equipped with a touch screen and retractable levers for operant tasks, along with a water port for reward delivery, all integrated into an individually housed home cage. This platform can be easily and inexpensively (∼$500 per unit) assembled using readily available materials or a 3D printer, following step-by-step instructions provided on our web site (https://operanthouse.sakura.ne.jp/index.html). As a result, this affordable platform enables automated operant conditioning tests on multiple mice in their home cages.

To demonstrate the effectiveness of the Operant House, we present two protocols for assessing working memory, a crucial cognitive function required for temporarily holding information or perception in attention to guide goal-directed behavior. Working memory deficits are observed in various neurological and neuropsychiatric disorders, including Alzheimer’s disease and schizophrenia. To evaluate working memory, we utilized a mouse model of Alzheimer’s disease (*App^NL-G-F^*), carrying APP mutations associated with familial Alzheimer’s disease without changing APP expression levels (Saito et al., 2014). Remarkably, the Operant House detected impaired working memory in *App^NL-G-F^* as early as 5 months of age, a finding that often goes undetected in conventional experiments (Sakakibara et al., 2018; Whyte et al., 2018; Latif-Hernandez et al., 2019). The Operant House will significantly contribute to the field of behavioral neuroscience and enhance our understanding of cognitive impairments in neurological and neuropsychiatric disorders.

## Material and methods

### Design of the Operant House

The design of the Operant House was based on conventional operant conditioning devices to ensure compatibility with previously established operant tasks. Mice were individually housed in a home cage measuring 215 x 145 x 205 mm. The cage was filled with finely minced wood chips (White Frakes, Oriental Yeast Company, Japan), prepared using a food processor, to serve as bedding. Mice had *ad libitum* access to food, but their access to water was restricted during the night to facilitate operant conditioning without inducing stress (Fig. 1A).

**Fig. 1.**
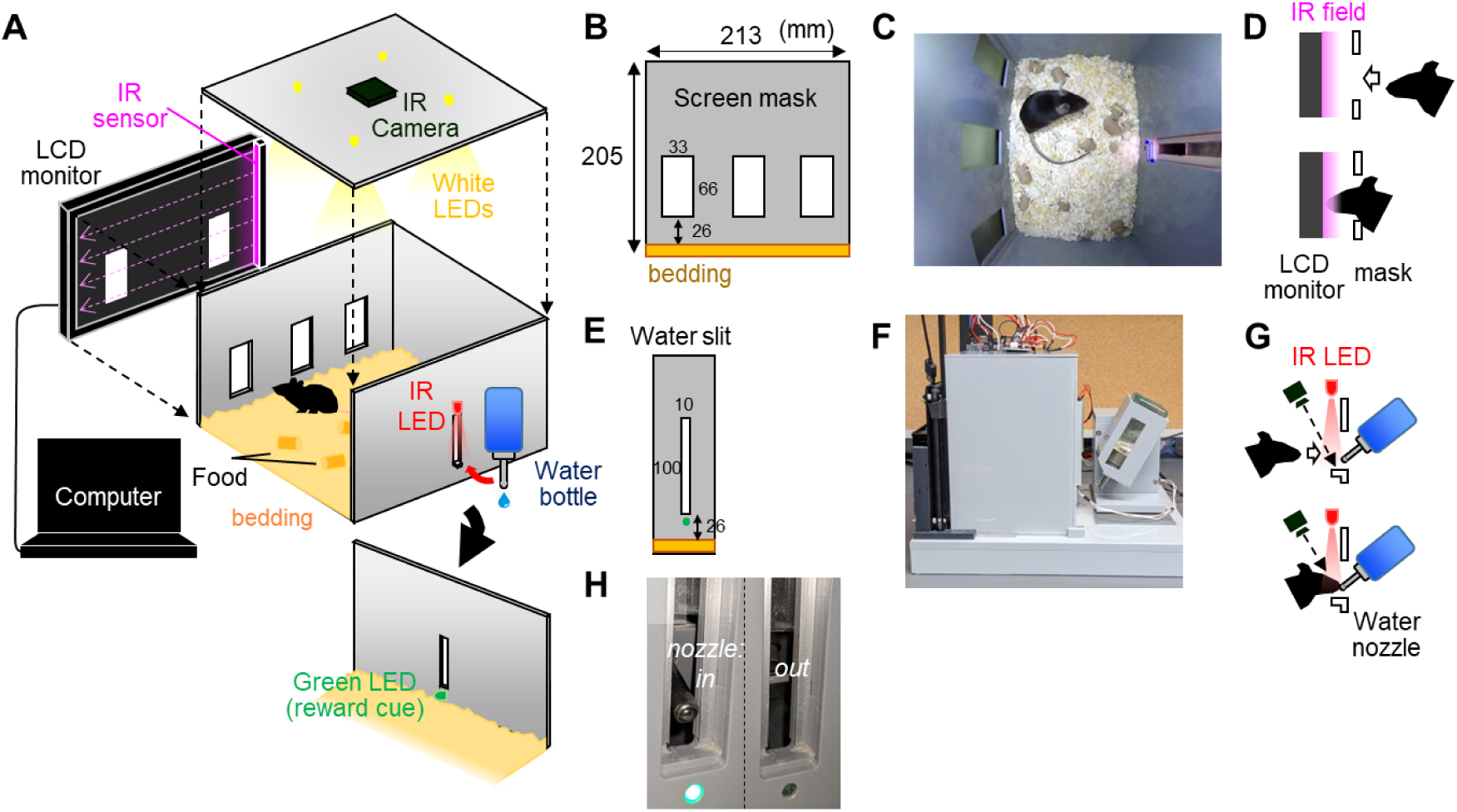
Assembly of the Operant House version 1. The front wall of the Operant House is composed of an LCD monitor (**A**) and a screen mask with open windows (**B**). Food is always available on the floor (**C**). An IR sensor generates an infrared field on the monitor’s surface to detect nose pokes (**D**). When the operant task is successful, a water bottle tilts forward through a gap in the rear wall (**E**), allowing the mouse to drink (**F**). During nighttime, an IR LED above the gap illuminates, and the drinking behavior of the mouse is recorded by an IR camera on the ceiling (**G**). A green LED below the gap lights up as a cue indicating a reward (**H**). White LEDs are installed on the ceiling, illuminating during the day and lighting temporarily at night as a punishment.

One of the chamber walls was equipped with a Liquid Crystal Display (LCD) monitor covered by a mask. A mouse could interact with the visual cue displayed on the LCD monitor through a slit in the mask (Fig. 1B-D). On the opposite side of the wall, a drinking water slit was provided, with a retractable water bottle (Fig. 1E, F). Upon successfully completing a trial of the operant task, a servo motor tilted the holder containing the water bottle. This action caused the mouth of the bottle to protrude through the slit into the chamber, allowing the mouse to access the water (Fig. 1G, H). Learning in the task was facilitated by activating a green Light Emitting Diode (LED) in the slot as a reward cue for correctly performing the operant task, while four white LEDs on the ceiling provided as an aversive stimulus which served as punishment (Fig. 1A). The same white LEDs were continuously turned on to illuminate the entire chamber during the daytime. The servo motor controlling the water bottle holder and the LEDs were operated by a microcontroller board (Arduino Uno, Arduino LLC, Italy) connected via a universal serial bus (USB). The mask of the LCD monitor was interchangeable, enabling the Operant House to perform various operant conditioning paradigms by combining different slit patterns in the mask with specific visual cues displayed on the LCD screen.

The mouse’s behavior was continuously recorded using an infrared (IR) camera (Fig. 1A). A Python 3 script controlled the operant tasks, outcomes, watering device, lighting, and IR camera. The open-source Python program, written in modules can be easily customized to suit the user’s needs with free integrated development environment (IDE) such as PyCharm community or Jupyter Notebook. The program featured a graphical user interface, ensuring user-friendliness. The system was compatible with remote-controlling software such as TeamViewer or Chrome Remote Desktop. Once all parameters were set on the first day, the entire experiment could run without requiring additional modifications.

A step-by-step instruction guide for constructing the Operant House, including the comprehensive information on the 3D printer, electronic parts, and Raspberry Pi 4, is available on the website (https://operanthouse.sakura.ne.jp/index.html).

#### Operant House Version 1

The chamber was constructed using polyvinyl chloride (PVC), cut by a workshop. A laptop computer was used to control devices connected to each chamber (Fig. 2A). An infrared sensor (AirBar, Sweden, or NNAMC3460PCEV, Neonode Inc.) positioned on an LCD monitor (connected via High-Definition Multimedia Interface [HDMI]) detected the mouse’s poking action through the mask panel (Fig. 1D).

**Fig. 2.**
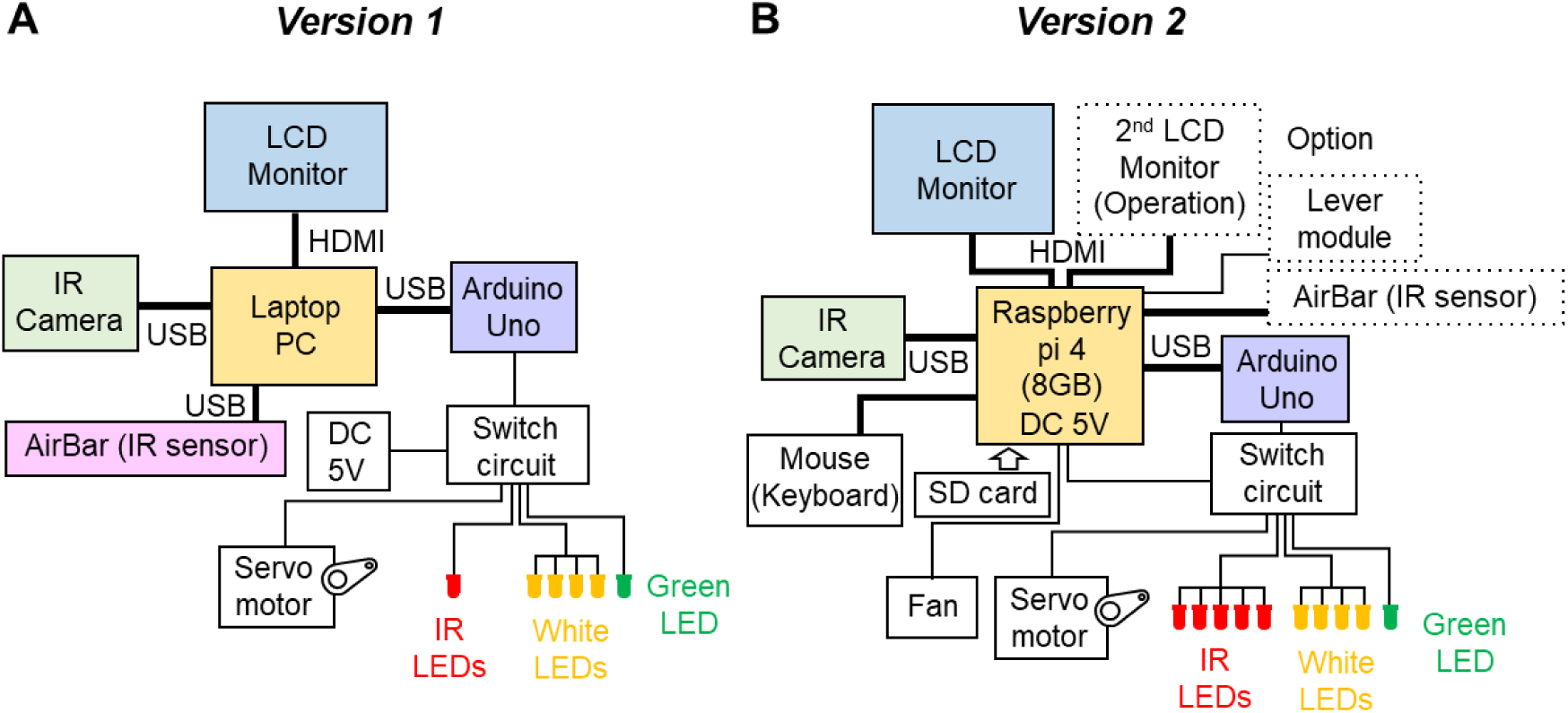
Block diagrams for the Operant Housse. To capture IR camera images and control peripheral devices, version 1 (**A**) uses a laptop PC, while version 2 (**B**) employs a single-board computer (Raspberry Pi 4). Additionally, to detect nose-poke actions, version 1 (**A**) uses a touchscreen equipped with an IR sensor (AirBar) placed on the LCD monitor. In contrast, version 2 (**B**) utilizes image recognition of mouse movements captured by an IR camera. Several optional devices are also available including operational monitor, lever modules and an AirBar.

#### Operant House Version 2

The chamber was redesigned to be compatible with a FDM 3D printer and polylactic acid filament (PLA) (Fig. 3A). A single-board computer (Raspberry Pi 4) and a Secure Digital (SD) card replaced the laptop computer (Fig. 2B). To reduce costs further, we replaced the infrared sensor with a motion-based detection system utilizing IR LEDs and an IR camera (Fig. 3B). This modification also allowed experimenters to adjust the detection sensitivity by altering the distance from the LCD monitor to the detection area (Fig. 3C). By combining these modifications, the Operant House Version 2 could be built for approximately $500 per unit.

**Fig. 3.**
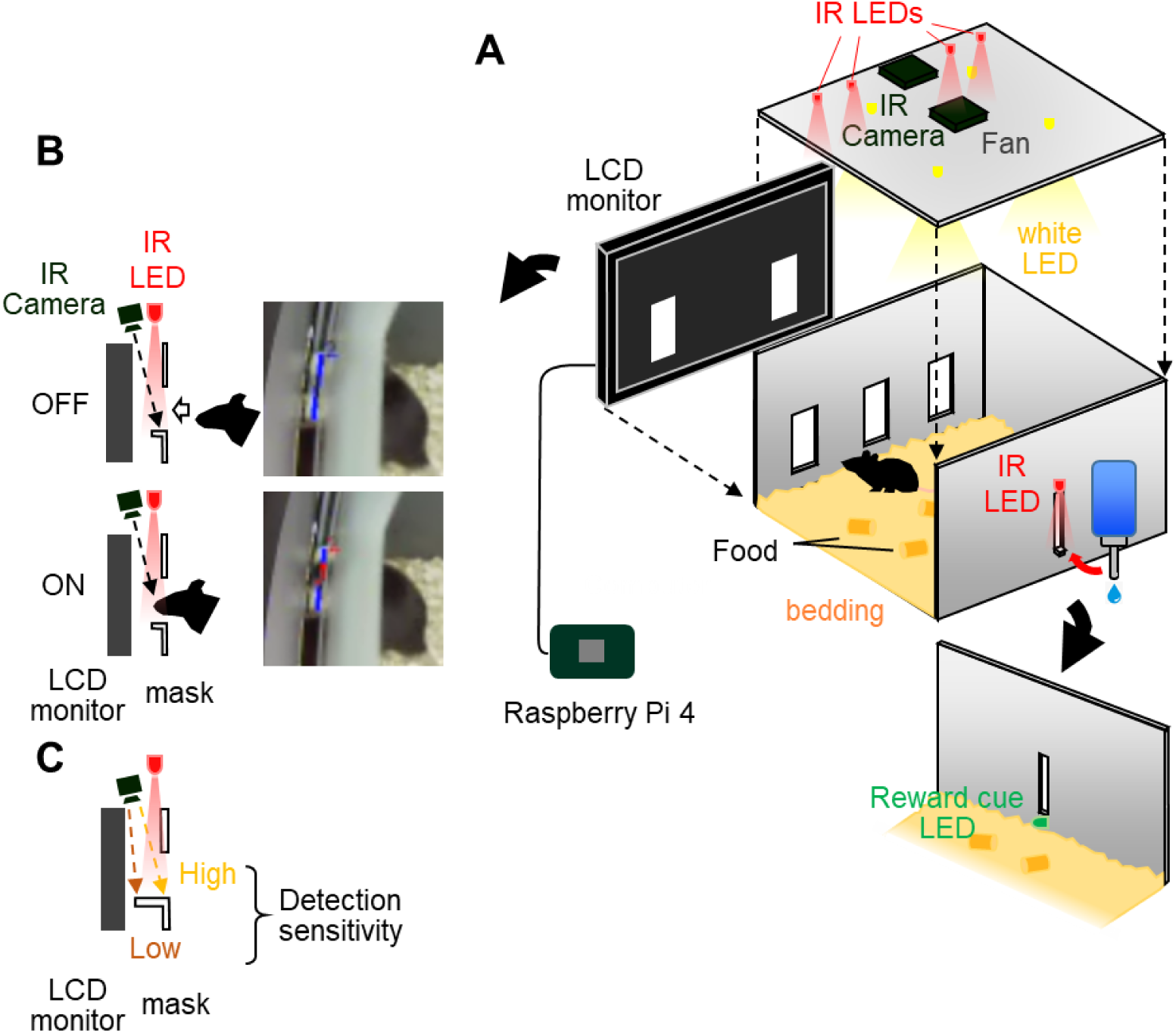
Assembly of Operant House version 2. Similar to Version 1, the Operant House comprises an LCD monitor and a screen mask with open windows, with food continuously available on the floor. Upon successful completion of the operant task, a water bottle tilts forward through a gap in the rear wall, enabling the mouse to drink (**A**). Unlike Version 1, nose-poke actions are detected using image recognition of mouse movements, eliminating the need for an IR sensor (**B**). This is achieved by adding four IR LEDs to the ceiling. The sensitivity of nose-poke detection can be fine-tuned by adjusting the angle of the IR camera (**C**). All devices are managed by a single-board computer, Raspberry Pi 4.

#### An Updatable Lever-Press Module

As an additional feature of Operant House Version 2, we introduced a modification that allowed the replacement of the wall featuring a drinking water slit with a wall containing a retractable lever switch (Supplementary Fig. S1A). The lever switch was designed to require minimal force for activation and would return to its original position automatically due to gravity (Supplementary Fig. S1B). To control the movement of the lever switch into and out of the chamber, a servo motor was incorporated (Supplementary Fig. S1C). This module significantly expanded the range of operant tasks that could be performed within the Operant House.

### Automated Operant Conditioning using the Operant House

On the first day of the tasks, mice were transferred to the Operant House chamber between 9:00-12:00 am for acclimation. The mice were only allowed access to water at night. The total duration of water access was continuously monitored to ensure adequate hydration of the mice. If the total water intake fell below the required level, additional access was provided after the end of the daily trials to maintain consistent water intake throughout the experiment. All the following tasks commenced at midnight (0:00), indicated by the illumination of a green LED and a brief 1-sec water supply.

### Delayed non-matching to position (DNMTP) test (Fig. 4A)

A mask with three apertures, each aligned with an illuminated rectangular panel on the LCD screen, was used to present visual cues. The protocol consisted of three blocks. Mice were advanced to the next block either after a predetermined number of days or when their performance reached a specified criterion level. Daily trials in each block were concluded when the total number of correct responses reached 80 or when 10 hours had elapsed since the start of the trials.

#### Block 1-1: VD1 task (Fig. 4B)

In each trial, one of the three panels was randomly illuminated. When the subject touched the illuminated panel, the panel light was turned off, the green LED at the water port was activated, and the subject was allowed access to water for 2 sec. The water nozzle retracted 2 sec after detecting the subject’s nose poke into the water slit. Touching an unilluminated aperture was considered an incorrect response, resulting in a 2-sec timeout. The inter-trial interval (ITI) was set at 5 sec. Since many subjects did not initially recognize the association between panels and the water reward, they were given 1-sec water access if they did not respond to the illuminated panels for longer than 6 min (stimulus water supply). VD1 was performed until the subject achieved 5 successive correct responses, at which point they were moved to visual discrimination task 2 (VD2).

#### Block 1-2: VD2 task (Fig. 4C)

VD2 differed from VD1 in two ways. Firstly, the stimulus water supply was not provided. Secondly, mice were punished with chamber light illumination (10 sec) when touching unilluminated panels. In the experiments involving ibotenic acid injection, VD2 was repeated for four days before the DNMTP pre-acquisition task. For experiments with the Alzheimer’s disease model, VD2 was repeated until the correct response exceeded 50%. The learning curves in control animals were similar in both conditions. In the analysis of wild-type animals using Operant House Version 2, VD2 was repeated until the correct response exceeded 65% for two consecutive days.

#### Block 2: DNMTP pre-acquisition task (Fig. 4D)

The DNMTP pre-acquisition task was designed to help mice recognize the rules employed in the DNMTP task. At the beginning of the sample phase, either the left or right panel flickered. When the mice touched the flickering panel, the panel turned off. After 1 sec, the middle panel illuminated, signaling the start of the delay phase. This sequence was repeated once more to indicate the end of the delay phase. The delay phase lasted 1 sec, followed by the choice phase. In the choice phase, the panel located opposite the sample panel was illuminated, and touching this panel led to access to a water reward for 2 sec. Touching the same panel that had been illuminated during the sample phase was considered an incorrect response and resulted in a 10-sec chamber illumination punishment. The ITI was 30 sec. The DNMTP pre-acquisition task was performed for one day in Version 1. In Operant House Version 2, the DNMTP pre-acquisition task was repeated until the correct rate exceeded 85% for two consecutive days.

#### Block 3: DNMTP task (Fig. 4E, F)

The DNMTP task was similar to the DNMTP pre-acquisition task, with the difference being that in the choice phase, both panels on the left and right sides were illuminated. The subject was required to remember the location of the sample panel and touch the correct panel in the choice phase. During the acquisition trials, the duration of the delay phase was set to 1 sec. The trials were repeated until the total success rate reached 80% (for *App^NL-G-F^* mice) or 75% (for experiments using Operant House Version 2) for two days. Most wild-type mice achieved this criterion within 1-2 weeks. Once the criterion was met, the duration of the delay phase was gradually increased (Fig. 4F). To prevent the mice from acclimating to a specific pattern of changes, two delay schedules, A and B, were randomly assigned to each trial. Trials using delay schedule B were analyzed for working memory.

**Fig. 4.**
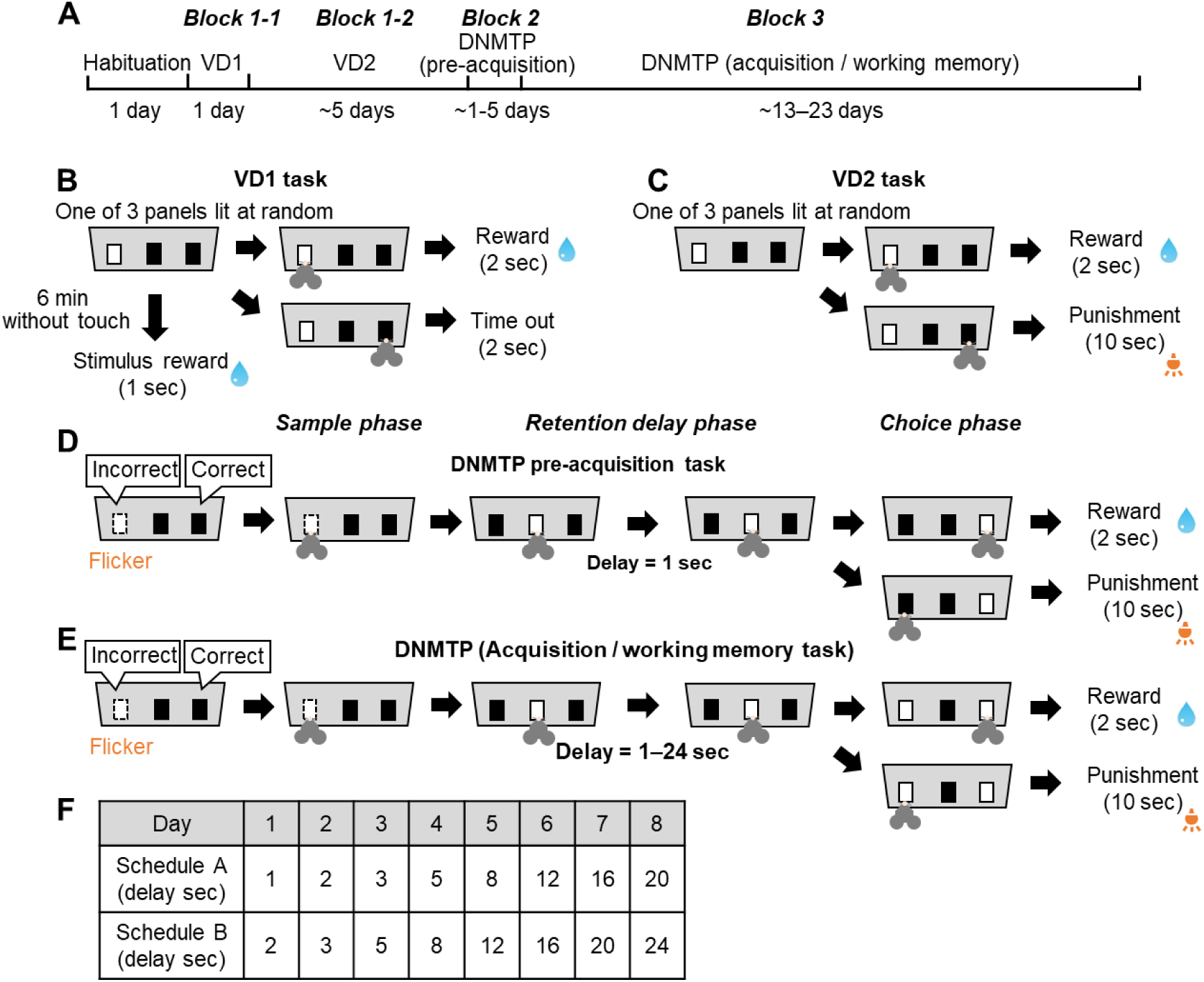
Protocol for delayed non-matching to position (DNMTP) test. (**A**) The DNMTP experiment consisted of three blocks: Block 1 (Visual Discrimination 1 [VD1] and VD2 tasks), Block 2 (DNMTP pre-acquisition task), and Block 3 (DNMTP task). (**B**) VD1 Task. Touching an illuminated panel was rewarded with water, while touching unilluminated panels resulted in a 2-sec timeout. If the subject did not respond to any panel within 6 min, they received a 1-sec water stimulus reward. (**C**) VD2 Task. Similar to VD1, except that incorrect responses were penalized with a 10-second light stimulus and the omission of the stimulus reward. (**D**) DNMTP Pre-Acquisition Task. Subjects were trained to complete a sequence of four touches to receive water. The first touch was on a blinking panel (left or right), the second and third touches were on the middle panel, and the fourth touch was on the panel opposite the initial touch. (**E**) DNMTP Task. The same as the pre-acquisition task, except during the choice phase, both left and right panels were illuminated. The interval between the second and third touches (on the middle panel) constituted the delay phase, starting at 1 sec until the criterion was met. (**F**) Delay Settings in the DNMTP Task. Two delay schedules (A or B) were applied randomly to each trial. Delay durations progressively increased over days.

### Two-choice spatial discrimination (TCSD) test

A mask with five apertures, each corresponding to illuminated rectangles on the LCD screen, was used to present visual cues. In the TCSD test, mice were required to memorize the correct position of the illuminated panel from the previous trial to receive a water reward. The experiment consisted of two blocks (Fig. 7A). In the first block, mice were trained to associate touching the illuminated panels with the water reward by performing visual discrimination tasks (VD1 and VD2). The ITI was set at 5 sec. Daily trials were terminated either when the total number of correct responses reached 150 or when 6 hours had elapsed since the start of the trials.

#### Block 1-1: VD1 task (Fig. 7B)

The VD1 task was conducted similarly to that of the DNMTP test (Fig. 4*B*), with the exception that one of the five panels was randomly illuminated. The stimulus water supply was also employed. Once the mice achieved five successive correct responses, they proceeded to the VD2 task.

#### Block 1-2: VD2 task (Fig. 7C)

The VD2 task was also identical to the VD2 in DNMTP test, except that five panels were used. The VD2 task was repeated until the correct response rate exceeded 50%.

#### Block 2: TCSD task (Fig. 7D)

In the TCSD task, two panels were simultaneously illuminated, and one of the two panels was randomly assigned as the correct choice at the beginning of trials. When the subject touched the correct panel, it was rewarded with 2 sec of water access. The incorrect choice was punished with chamber illumination for 5 sec. The position of the correct choice was reversed to the opposite side when the subject achieved more than 7 correct responses in the most recent 8 trials. The TCSD task was repeated for 3 days.

### Lever-pressing conditioning test (Supplementary Fig. S1)

Following a 24-hour water deprivation period, the subjects were placed in the chamber at 9:00 pm, and the task commenced immediately. To begin trials, both right and left levers were inserted into the chamber and pressing either lever resulted in a reward of 2-sec access to water. Daily trials were terminated when the subject pressed the levers 80 times, which was followed by the retraction of the levers. A wire grid wall was inserted in the middle of the chamber on the first day to reduce the chamber space and prompt the mice to press the levers. The wire grid wall was removed on the second day.

### Animals

All procedures regarding animal care and treatment were conducted in accordance with the guidelines established by the Animal Resource Committee of Keio University and the RIKEN Center for Brain Science.

Homozygous *App^NL-G-F^* knock-in mice were obtained from RIKEN Bioresource Center (#RBRC06344) and crossed with C57BL/6J (Crlj) to produce heterozygous breeding pairs. The resulting mice were used to obtain mice homozygous for *App^NL-G-F^*, while wild-type littermates were used as control subjects. The littermates were housed in the same cages, maintained under a 12:12 hour light/dark cycle, at a temperature of 22 ± 1°C, and provided with ad libitum access to food and water. For the DNMTP experiment, both female and male mice at the age of 5 months were used. In the TCSD experiment, we used 5-month-old male mice. Wild-type mice at the age of 4 months were used for the lever-pressing conditioning test. Histological and electrophysiological analyses were conducted using male mice.

### Ibotenic acid injection

Male wild-type mice (C57BL/6J, Japan SLC) at the age of 2 months were subjected to an intraperitoneal injection of a mixed anesthetic comprising medetomidine (0.3 mg/kg), midazolam (4 mg/kg), and butorphanol (5 mg/kg). Ibotenic acid (Abcam, ab120041) was prepared as a solution at a concentration of 10 mg/ml in saline and stored at −20°C until use. To verify the accuracy of the injection site, Red Retrobeads IX (LumaFluor) were added at a volume ratio of 1:20 to the ibotenic acid solution. A total of 400 nl of the resulting mixture was injected into the bilateral prefrontal cortex using a stereotaxic device (Stoelting) and Nanoliter 200 (World Precision Instrument). The injection coordinates were as follows: anterior-posterior (AP) +2.75 mm, medial-lateral (ML) 0.4 mm, and depth from the pia 0.8 mm. For the control group, saline and Red Retrobeads IX were injected. Following the surgery, the mice were allowed to recover for a period of five days before being subjected to the DNMTP task.

### Immunostaining of c-Fos after the DNMTP task

Female wild-type mice (C57BL/6J, Japan SLC) aged 2-3 months were trained in the DNMTP task following the protocol described above. On the day when the mice performed the task with a delay of 8-12 sec, they were perfused with 4% paraformaldehyde (PFA) in phosphate-buffered saline (PBS) 60-90 min after the task began. Control mice were housed individually and were also perfused on the same day. The brains were post-fixed overnight with 4% PFA, and 100 μm sections were obtained using a vibratome (DTK-1000, Dosaka).

The brain slices were treated with a blocking solution containing 2% donkey serum and 0.3% Triton-X, and then incubated with an anti-c-Fos antibody (mouse monoclonal [2H2], Abcam, ab208942, 1:1000) dissolved in the same blocking solution. After three washes with PBS, the sections were incubated with the secondary antibody (donkey anti-mouse Alexa 555, Invitrogen, 1:500) and Neurotrace 600/647 (ThermoFisher, 1:500) for three hours at room temperature. Following three additional washes with PBS, the sections were mounted in 70% glycerol, and images were captured using a 10 x objective lens on a confocal microscope (FV1000, Olympus). Projection images from five z-sections were analyzed using Fiji software. Images were first filtered with a median filter, and the density of c-Fos positive cells was quantified. Cells exhibiting c-Fos intensity greater than three times the background were defined as c-Fos positive cells.

### Immunostaining of Aβ and synaptic markers

Acute coronal slices of the medial prefrontal cortex (mPFC) with a thickness of 300 µm were obtained following the same procedure described in the electrophysiology section below. The slices were fixed in 4% PFA in PBS overnight at 4 °C. Cryoprotection was achieved by immersing the slices in a 30% sucrose solution in PBS, followed by embedding them in Tissue-Tek embedding medium (Electron Microscopy Sciences, Hatfield, PA, USA). The frozen samples were then cut into 20 μm-thick coronal sections using a cryostat (CM1850, Leica).

Sections were treated with a blocking solution consisting of 10% donkey serum and 0.1% Triton-X in PBS for 30 min and incubated overnight with primary antibodies in 0.1% Triton-X / PBS. The following primary antibodies were used: anti-VGluT1 (rabbit, Frontier Institute, 1:500), anti-VGAT (guinea pig, Frontier Institute, 1:1000), anti-Iba1 (rabbit, Sigma, 1:500), and anti-Aβ antibody (mouse monoclonal [82E1], IBL, 1:500). After three washes with 0.1% Triton-X / PBS, the sections were incubated for three hours at room temperature with the appropriate secondary antibodies (goat anti-mouse IgG1 Alexa 488, goat anti-mouse IgG1 Alexa 647, donkey anti-rabbit Cy3, or donkey anti-guinea pig Alexa 488, Jackson, 1:500) and Neurotrace 600/647 (ThermoFisher, 1:500). Following three additional washes with 0.1% Triton-X / PBS, the sections were mounted in Vectashield Mounting Medium (Vector Laboratories). Images were captured using either a confocal microscope (FV1000, Olympus) with a 4x objective lens or an Olympus SR-OSR microscope with either a 10 x or 100 x objective lens. For images obtained using the 100x objective lens, z-projected images from 23 z-sections were acquired and analyzed using Fiji software. The analysis was performed semi-automatically using a custom macro. Background noise was reduced by applying background subtraction using the rolling ball method (5-50 pixels). Regions of interest corresponding to Aβ plaques were determined based on intensity using Otsu binarization and size thresholding (above 50 μm^2^). The mean signal intensity outside the plaques was obtained from three different positions within areas of 370 μm^2^.

### Electrophysiological recordings in acute cortical slices

Coronal slices of the mPFC with a thickness of 300 µm were prepared from 5-6-month-old mice. The mice were anesthetized with isoflurane, and acute brain slices were obtained using a vibratome (DSK-1000N, Leica) in chilled (0-4 °C) normal artificial cerebrospinal fluid (ACSF) containing (in mM): 126 NaCl, 3 KCl, 1.3 MgSO4, 2.4 CaCl_2_, 1.2 NaH_2_PO_4_, 26 NaHCO_3_, and 10 glucose. The slices were maintained at a temperature of 34 °C with 95% O_2_ and 5% CO_2_ during the cutting procedure. Before recording, the slices were incubated for a minimum of 30 min in normal ACSF at 34 °C, which was continuously bubbled with 95% O_2_ and 5% CO_2_.

Pyramidal neurons were identified based on their characteristic pyramidal soma shape using infrared differential interference contrast optics (BX51, Olympus) and a ×60 water immersion objective lens (LUMPlanFI/IR, NA 0.9, Fig. 10A) along with a CCD camera (Cool SNAP Kino, Teledyne Imaging). Whole-cell patch-clamp recordings were performed on neurons in layer 2/3 and layer 5 of the mPFC, including the anterior cingulate, prelimbic, and infralimbic areas, at room temperature. The patch pipettes had a resistance of 4-6 MΩ and were filled with an intracellular solution consisting (in mM): 130 K-gluconate, 8 KCl, 1 MgCl_2_, 0.6 EGTA, 10 HEPES, 3 MgATP, 0.5 Na_2_GTP, and 10 Na-phosphocreatine (pH 7.3 with KOH).

To assess the intrinsic membrane properties and excitability of pyramidal neurons, 500-ms step currents of varying amplitudes were injected in current-clamp mode. Current injection experiments were performed within 5 min of establishing the whole-cell patch-clamp configuration.

For measuring evoked EPSCs/IPSCs and miniature EPSCs/IPSCs, a pipette solution containing (mM) 130 Cs-gluconate, 6 KCl, 10 NaCl, 10 HEPES, 0.16 CaCl_2_, 2 MgCl_2_, 0.5 EGTA, 4 Na-ATP, and 0.4 Na-GTP (pH 7.3, adjusted with CsOH) was used. EPSCs and IPSCs were recorded at holding potentials of −70 mV and 0 mV, respectively. To assess the inhibitory/excitatory (I/E) balance, EPSCs and IPSCs were evoked by electrical stimulation using a pair of bipolar tungsten electrodes (FHC, Corporation), positioned near the recorded neuron (Fig. 10A, inter-stimulus interval = 20 s). Ten to twenty consecutive traces were averaged to obtain the mean EPSCs or IPSCs. The baseline current was defined as the average current in a 20-ms window prior to the electrical stimulation, and the peak EPSC and IPSC were determined by calculating the absolute value of the negative or positive peak current minus the baseline. To measure mEPSCs and mIPSCs, recordings were conducted in the presence of 1 μM TTX. The experimental setup included a Multiclamp 700B amplifier, Digidata 1440A converter, and pClamp10 software (Molecular Devices). Data were digitized at a sampling rate of 20 kHz, filtered at 4 kHz, and analyzed using custom-made MATLAB programs (MathWorks). Cells with a seal resistance between 1 GΩ and 35 MΩ were included in the analysis.

### Electrophysiological recordings in acute hippocampal slices

Hippocampal slices with a thickness of 300 μm were prepared from 5-month-old *App^NL-G-F^* mice and littermate wild-type mice. The mice were decapitated under isoflurane anesthesia. For the dissection, a cutting solution composed of the following concentrations (in mM) was used: 229 mannitol, 3 KCl, 23 NaHCO_3_, 1.23 NaHPO_4_, 0.5 CaCl_2_, 3 MgCl_2_, and 10 glucose, with a pH of 7.4. The cutting solution was continuously bubbled with a mixed gas of 95% O_2_ and 5% CO_2_. After the dissection, the slices were kept at room temperature in ACSF solution with the following concentrations (in mM): 124 NaCl, 3 KCl, 2 CaCl_2_, 2 MgCl_2_, 1.23 NaH_2_PO_4_, 26 NaHCO_3_, and 10 glucose, with a pH of 7.4. The ACSF solution was also bubbled with a mixed gas of 95% O_2_ and 5% CO_2_.

Field excitatory postsynaptic potentials (fEPSPs) were recorded in the stratum radiatum of the CA1 region using the MED64 system (Alpha MED Scientific). The hippocampal slices were placed on the MED64 probe, which consisted of an array of 64 planar microelectrodes arranged in an 8 x 8 pattern (P515A, Alpha Med Scientific). The recording probe covered the entire hippocampal CA1 region. During the experiments, the slices were continuously perfused with ACSF at a rate of 1.5 ml/min at room temperature. Monopolar, biphasic constant current pulses with a duration of 0.1 msec were applied to the Schaffer collateral pathway at 30-sec intervals for electrical stimulation. The amplitude of the electrical stimulation was determined based on the input-output curve, and we used 1/3 of the amplitude that induced the maximum fEPSP response.

To induce long-term potentiation (LTP), a stable baseline of fEPSP amplitude was maintained for at least 10 min, followed by high-frequency stimulation consisting of 100 Hz and 100 pulses. The data were analyzed using the data acquisition software Mobius (Alpha-Med Scientific).

### Data analysis and statistics

All results are presented as means ±SEM. Two-way repeated-measures ANOVA was performed followed by Bonferroni *post hoc* tests, unless otherwise stated. To identify outliers, all datasets underwent the Smirnov-Grubbs outlier test at a significant level of 0.05. Statistical significance was defined as p < 0.05.

## Results

### Assessment of working memory through the DNMTP task using Operant House

We developed Operant House to automatically perform operant conditioning tasks in individually housed home cages (Figures 1–3, refer to Methods for detailed description). To validate its effectiveness, we designed a protocol for evaluating working memory using the delayed non-matching to position (DNMTP) test (Fig. 4). In this test, the subject was required to remember the position of one of three illuminated panels during the sample phase and to choose the panel on the opposite side during the choice phase to receive a water reward. The protocol consisted of three blocks (Fig. 4A, refer to Methods for detailed description):

In the first block, mice performed visual discrimination tasks, VD1 and VD2, and learned to touch illuminated panels to receive water rewards (Fig. 4B, C). By the end of day 1, the average correct rate for the VD1 task in wild-type mice was significantly higher than the chance level (Fig. 5A, *p* = 0.012 by one sample t-test). Throughout the VD2 task, the success rate gradually improved, reaching 76.5% on the fourth day, significantly higher than the first day (Fig. 5B, *p* = 0.037 by one-way repeated ANOVA with Bonferroni).

**Fig. 5.**
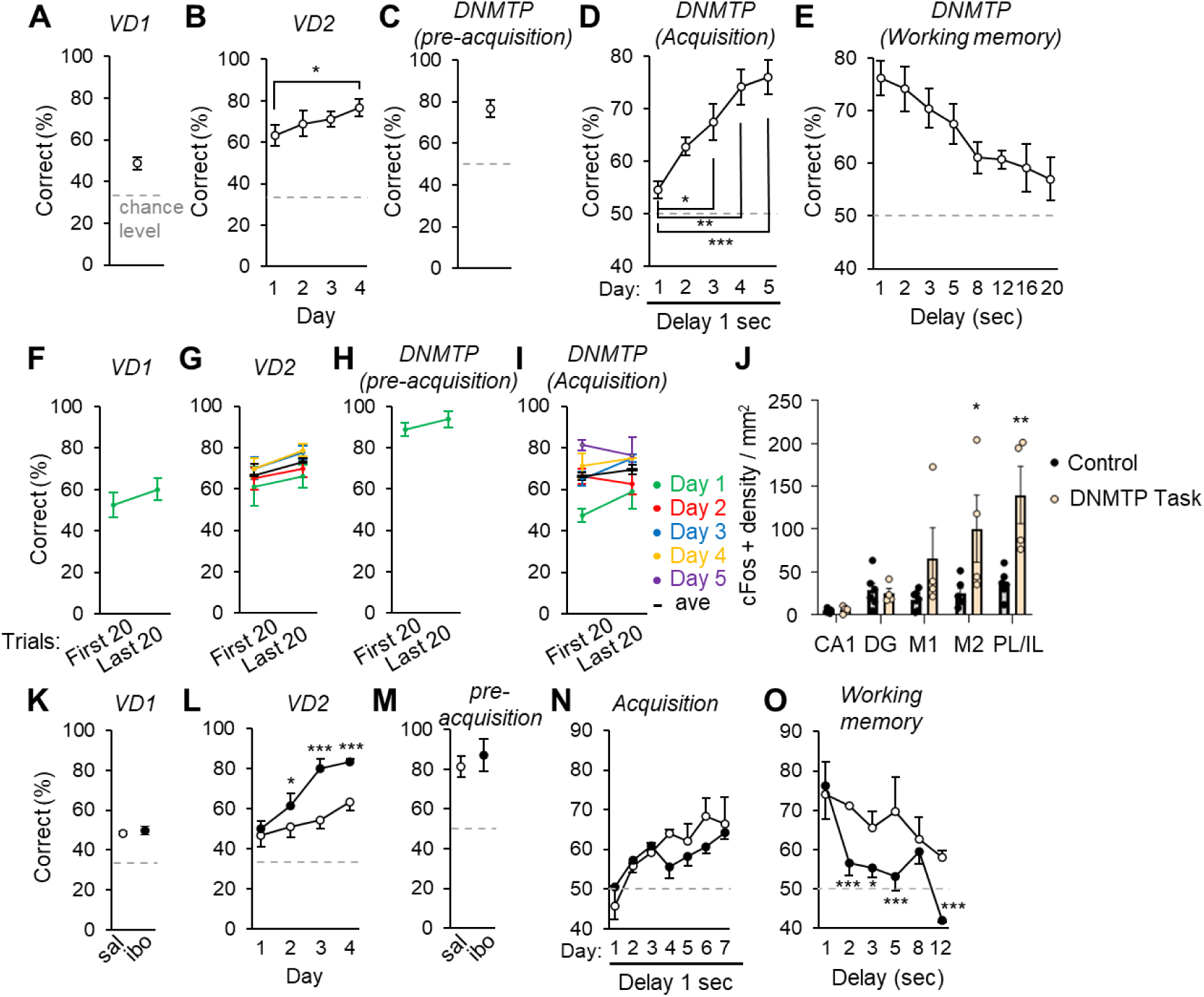
Working memory assessment using the DNMTP test in wild-type mice. (**A-E**) The average correct rates were calculated across all trials per day for the VD1 (**A**), VD2 (**B**), DNMTP pre-acquisition (**C**), DNMTP acquisition with a 1-second delay (**D**), and DNMTP working memory tasks (**E**). The dotted lines indicate chance-level performance. Note the decline in correct rates as delay intervals increased in the working memory task (**E**). (**F-I**) Comparison of correct rates between the first and last 20 trials of each task on the same day. No significant differences were observed for VD1 (**F**, paired t-test: *p =* 0.495), VD2 (**G**, two-way repeated ANOVA: F1,3 = 1.199; *p =* 0.315), DNMTP pre-acquisition (**H**, paired t-test: *p =* 0.423), or DNMTP acquisition tasks (**I**, two-way repeated ANOVA: F1,3=1.222; *p =* 0.311). (**J**) c-Fos activity. A significant increase in c-Fos expression was observed in M2 and PL/IL regions of mice performing the DNMTP working memory task. The density of c-Fos-positive neurons was measured in brains fixed 60–90 min after initiating the task (with delays of 8-12 sec). (**K-O**) Average correct rates for VD1 (**K**), VD2 (**L**), DNMTP pre-acquisition (**M**), DNMTP acquisition with a 1-sec delay (**N**), and DNMTP working memory tasks (**O**) in mice injected with saline (unfilled circles) or ibotenic acid (filled circles). *n* = 4 mice (**A-I**); *n* = 6 and 4 mice for control and NDMTP task group, respectively (**J**); *n* = 3 mice (**K-O**) per group. One-way repeated ANOVA with Bonferroni post-hoc tests: \*\*\**p* < 0.0001; \*\**p* <0.001; \**p* <0.01 (**B, D, E**). Two-way repeated ANOVA with Bonferroni post-hoc tests: \*\*\**p* < 0.0001; \*\**p* < 0.001; *p < 0.05 (**J, L, N, O**). All graphs show mean ± SEM.

In the second block, mice performed the DNMTP pre-acquisition task. Mice received a water reward after touching the correct panel during the choice phase. The correct panel was on the opposite side of the panel presented during the sample phase. The choice phase followed the sample phase after a fixed 1-sec delay phase. In this DNMTP pre-acquisition task, the correct panel was explicitly illuminated during the choice phase to help the mice recognize the rules of the DNMTP task (Fig. 4D). The success rate increased daily in wild-type mice, reaching 76.5% within 5 days (Fig. 5D, *p* = 0.008 vs. 1st day by one sample t-test).

In the third block, working memory was assessed by increasing the duration of the delay phase (Fig. 4E) according to a set schedule (Fig. 4F). The correct response rate decreased with longer delay times (Fig. 5E), but even with a 20-second delay, the correct rate remained significantly higher than chance level (One sample t-test: *p =* 0.039), indicating that the mice could retain positional memory for 20 sec (Fig. 5E).

Motivation is a critical factor influencing behavioral outcomes in tasks. Because the level of thirst influences motivation in our tasks, the correct rate and the frequency of trials may decline with reduced motivation after repeating trials and obtaining rewards. However, we did not observe a decrease in correct rates as trials were repeated each day. The correct rates in the first and last 20 trials were similar in VD1, VD2, DNMTP pre-acquisition, and DNMTP acquisition tasks (Fig. 5F-I). This trend was reflected in the average correct rate for 10-trial bins each day (Supplementary Fig. S2A-E). Most mice completed trials after achieving 80 correct responses (Supplementary Fig. S2F-J). The average duration of the entire trials per day was less than 3 hours except for the DNMTP task, which included delay phase between sample and choice phase (Supplementary Fig. S2K-O). The duration of each trial in the VD2 and DNMTP acquisition tasks was similar across repeated trials in a day (One-way repeated ANOVA: VD2, *F* = 1.731, *p =* 0.230; DNMTP acquisition, *F* = 1.216; *p =* 0.355) (Supplementary Fig. S2Q, S). These findings suggest that motivation remained constant despite repeated trials throughout the experiment.

To identify the brain regions responsible for the DNMTP task, we performed immunostaining for c-Fos, a marker of neuronal activation. We observed significantly increased c-Fos immunoreactivity in the higher motor area (M2) and the prelimbic and infralimbic cortex (PL/IL) of mice that underwent the DNMTP task compared to homecage control mice (Fig. 5J).

To further explore the role of these c-Fos-positive brain regions in the DNMTP task, we lesioned bilateral PL/IL and M2 regions through injection of ibotenic acid (IBO). Both IBO- and saline-injected mice displayed similar correct response rates in each block with repeated trials (Supplementary Fig. S3A-D). Additionally, both IBO- and saline-injected mice completed trials similarly by achieving 80 correct responses (Supplementary Fig. S3E-H), indicating that motivation remained unaffected by IBO treatment. While IBO-treated mice performed well in most tasks in Block 1-3 (Fig. 5K, M, N; Supplementary Fig. S3I, K, L, M, O, P), they exhibited significantly lower correct rates than controls in the DNMTP task as the delay time increased (Fig. 5O, *p* = 0.0412 by ANOVA). The DNMTP task can be conducted using Operant House Version 2 with a slight modification in the protocol (Supplementary Fig. S4A-F), as described in the Methods section. These results suggest that the DNMTP task effectively assesses working memory in wild-type mice, which depends on the activity of PI/IL and/or M2.

### The DNMTP task detected early working memory deficit in female *App^NL-G-F^* mice

In humans, a working memory deficit is suggested to be one of the earliest signs of pre-symptomatic Alzheimer’s disease (AD) (Kirova et al., 2015; Han et al., 2017). To test the effectiveness of the Operant House, *App^NL-G-F^* mice, which harbor mutations found in familial AD patients, were subjected to the DNMTP test. The results from VD1 showed that the correct rate of female *App^NL-G-F^* mice was higher than controls (Student’s t-test, *p =* 0.041) (Fig. 6A). No change was observed in VD2 and the DNMTP pre-acquisition task (VD2, genotype: *F*_1,10_ = 0.882, *p =* 0.370; DNMTP pre-acquisition, Student’s t-test: *p =* 0.579) (Fig. 6B, C). However, the performance in DNMTP acquisition task differed significantly between wild-type and female *App^NL-G-F^* mice. While wild-type mice reached the criteria of the DNMTP acquisition task (success rate > 80% for 2 successive days with 1-sec fixed delay) in 13 ± 1 days (mean ± sem), female *App^NL-G-F^* homozygous mice took 23 ± 3 days, significantly longer than the wild-type (Student’s t-test: *p =* 0.016) (Fig. 6D, E). After reaching the criteria above, the mice performed DNMTP task with a longer delay time to assess working memory. The correct rate in female *App^NL-G-F^*mice showed a significant reduction compared to wild-types when the delay time was longer than 2 sec (genotype: *F*_1,10_ = 18.106, *p =* 0.0017) (Fig. 6F). Consequently, the number of incorrect responses and the number of total trials per day was significantly higher in *App^NL-G-F^* mice (incorrect: *F*_1,10_ = 59.118, *p* < 0.0001; total: *F*_1,10_ = 13.465, *p* = 0.0043) (Fig. 6H). To assess the impact of motivation levels, we examined the temporal variation of correct responses over trials within a day. Neither wild-type nor *App^NL-G-F^* mice showed any difference in the correct rate with repeated trials (Supplementary Fig. S5A-D). In addition, no difference was observed in the total number of trials or the average time spent per trial (Supplementary Fig. S5E-P). These results suggest female *App^NL-G-F^* mice had impaired working memory while motivation and basal activity were unaffected.

**Fig. 6.**
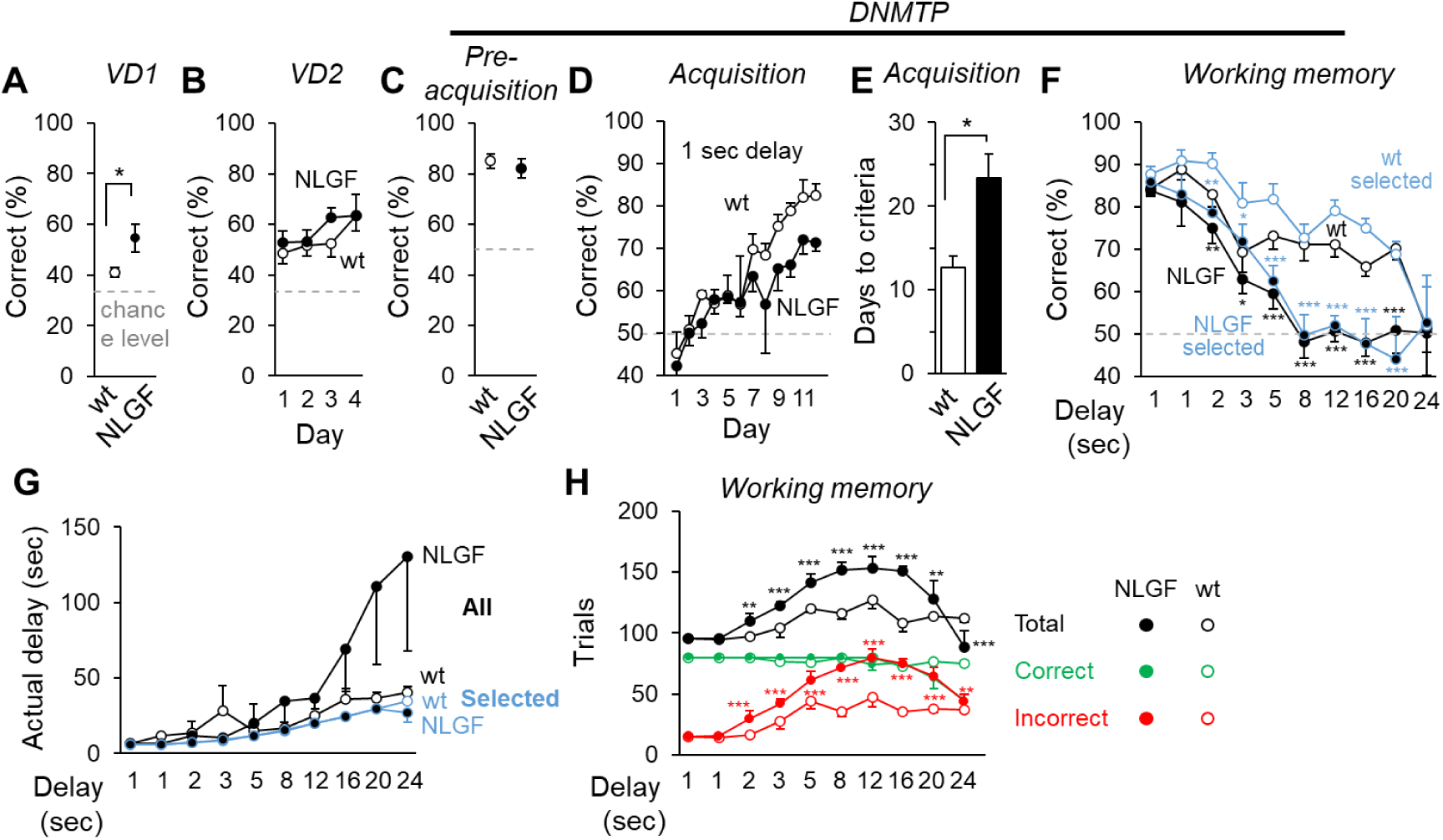
Female *App^NL−G−F^*mice exhibit working memory deficits in the DNMTP test at 5 months of age. (**A-C**) Average correct rates for VD1 (**A**), VD2 (**B**), and DNMTP pre-acquisition tasks (**C**) in wild-type (open circles) and *App^NL−G−F^*(filled circles) mice. Dotted lines indicate chance-level performance. (**D-E**) Daily improvements in correct rates during the DNMTP acquisition task with a 1-sec delay were slower in *App^NL−G−F^*mice (**D**). These mice required more days to reach the performance criterion (80% correct for two consecutive days) (**E**). (**F**) In the DNMTP working memory task, correct rates in *App^NL−G−F^* mice dropped sharply when the delay was 2 sec or longer. Blue-outlined plots represent trials meeting the criteria for actual delay times (refer to the main text). (**G**) No difference in actual delay times was observed between wild-type and *App^NL−G−F^* mice when comparing trials that met the criteria for actual delay time (Blue-outlined plots). (**H**) *App^NL−G−F^* mice performed more total trials in the DNMTP working memory task. This increase was observed in incorrect trials but not in correct trials. Student’s t-test: \**p* < 0.05 (**A, E**). Two-way repeated ANOVA with Bonferroni post-hoc tests: \*\*\**p* < 0.0001 (**F, H**). *n* = 6 mice per genotype.

In the DNMTP task, mice received a reward for touching the correct panel after the delay phase, which started and ended with the illumination of the middle panel. Consequently, the actual delay time between the sample and the choice phase, varied across trials. For instance, in some trials, mice were observed eating or sleeping during the delay phase, leading to longer actual delay time. Although not statistically significant (genotype: *F*_1,10_ = 1.117, *p =* 0.315), *App^NL-G-F^*mice tended to have longer actual delay times compared to wild-type mice (Fig. 6G). To exclude the potential confound by the variation in actual delay time, we selected trials whose actual delay time was within 1.3 times the specified delay time plus 10 seconds. These selected trials accounted for approximately two-thirds of all trials, and the average delay time was comparable between *App^NL-G-F^* and wild-type mice (Fig. 6G, blue outlines). Nevertheless, the percentage of correct responses in these trials remained significantly lower in *App^NL-G-F^* mice (genotype: *F*_1,10_ = 15.327, *p =* 0.0029) (Fig. 6F, selected trials), a result confirming impaired working memory.

Interestingly, male *App^NL-G-F^* mice did not exhibit any differences in the VD1/2 or DNMTP tasks (Student’s t-test: VD1, *p =* 0.213; DNMTP pre-acquisition, *p =* 0.411; two-way repeated ANOVA, genotype effects: VD2, *F*_1,9_ = 2.238, *p =* 0.169; DNMTP acquisition, *F*_1,9_ = 0.043, *p =* 0.840; DNMTP working memory, all trials, *F*_1,9_ = 0.527, *p =* 0.486; selected trials, F_1,9_ = 0.007, *p =* 0.934) (Supplementary Fig. S6). Taken together, these results indicate that sex-dependent working memory deficit in female *App^NL-G-F^* mice at early stages (Kundu et al., 2021) could be detected by the DNMTP task in the Operant House.

### The TCSD test detected impaired working memory in male *App^NL-G-F^* mice

To assess working memory by an independent method and to detect any changes in male *App^NL-G-F^* mice, we established a two-choice spatial discrimination (TCSD) test, in which mice were required to memorize the correct panel from two options by recalling the position from the previous trial (Fig. 7, refer to Methods for detailed description). The correct position was switched to the opposite side after the mice achieved more than 7 correct responses in the most recent 8 trials. Male *App^NL-G-F^* mice and their wild-type littermates at 5 months old were subjected to the TCSD task. In block1, VD1 and VD2 tasks, in which mice needed to touch a single lit panel, no significant differences were observed between genotypes (Student’s t-test: VD1, *p =* 0.271; VD2, *p =* 0.116) (Fig. 8A, B). In block 2, the TCSD test revealed lower correct rates in male *App^NL-G-F^* mice over 3 days (Fig. 8C; *F*_1,12_ = 33.685, *p* < 0.0001) and in each 50-trial bin (Fig. 8D; *F*_1,12_ = 15.497, *p =* 0.002). *App^NL-G-F^* mice also exhibited a significant reduction in the number of reversals (Fig. 8E; *F*_1,12_ = 19.056, *p =* 0.0009), with an increased average trial number per reversal (Fig. 8F; *F*_1,12_ = 8.513, *p =* 0.0129), while total trial numbers were unaffected (Fig. 8G; *F*_1,12_ = 3.210, *p =* 0.0984). These results suggest impaired working memory in male *App^NL-G-F^* mice.

**Fig. 7.**
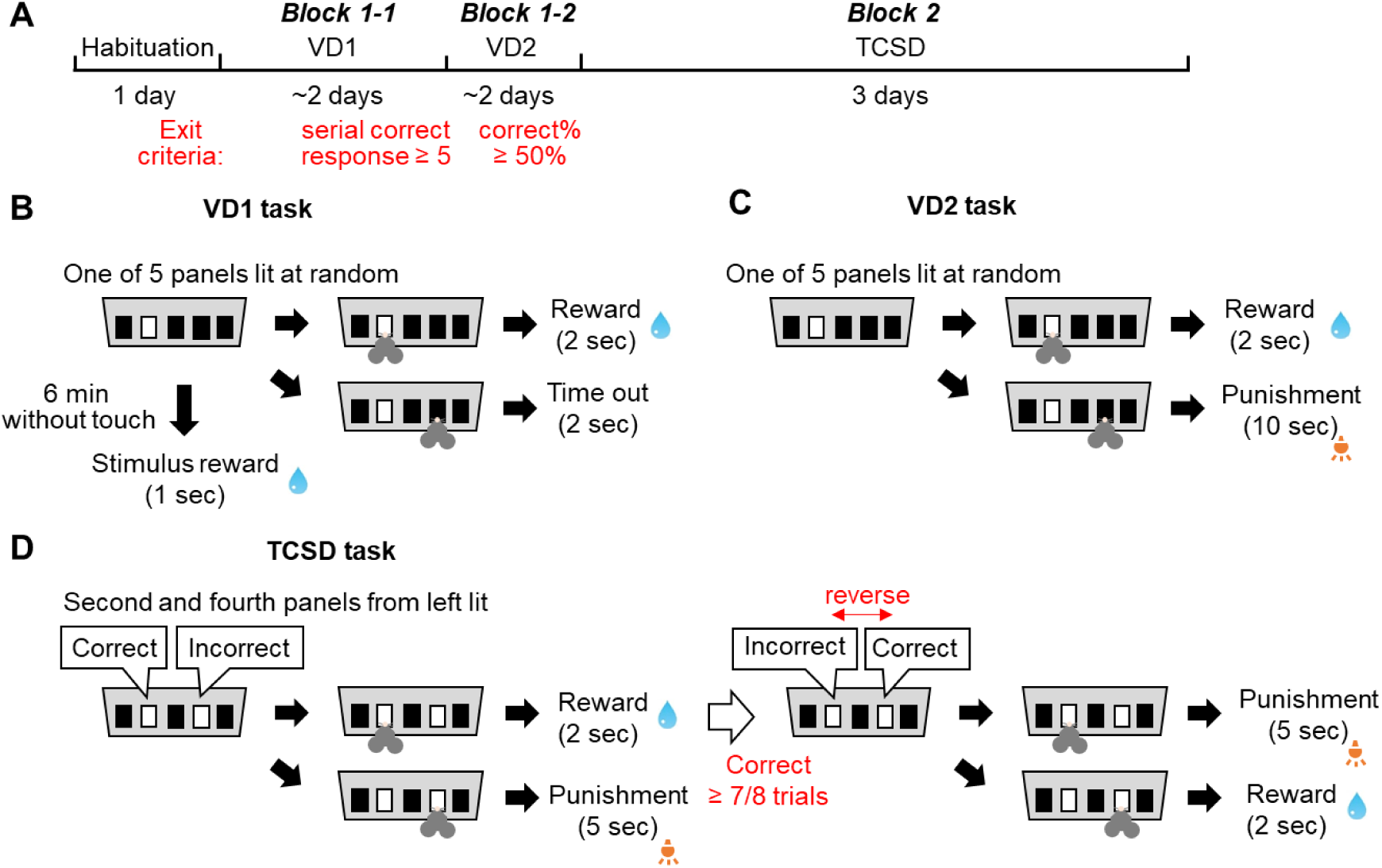
Protocol for two-choice spatial discrimination (TCSD) tests. (**A**) The TCSD test consisted of two blocks: Block 1 included Visual Discrimination 1 (VD1) and VD2 tasks, and Block 2 was the TCSD task. (**B**) VD1 Task. Touching the illuminated panel was rewarded with water, while touching unilluminated panels resulted in a 2-sec timeout. If the subject failed to respond within 6 min, a 1-sec water reward was provided as a stimulus. (**C**) VD2 Task. This task was similar to VD1, with the omission of stimulus rewards and the addition of a punishment for incorrect responses, consisting of a 10-sec light stimulus. (**D**) TCSD Task. At the start of the trials, the left or right panel was randomly designated as the correct choice. Selecting the correct panel resulted in a 2-sec reward, while choosing the incorrect panel triggered a 5-sec punishment. The correct panel switched after the subject achieved 7 correct responses out of the last 8 trials.

**Fig. 8.**
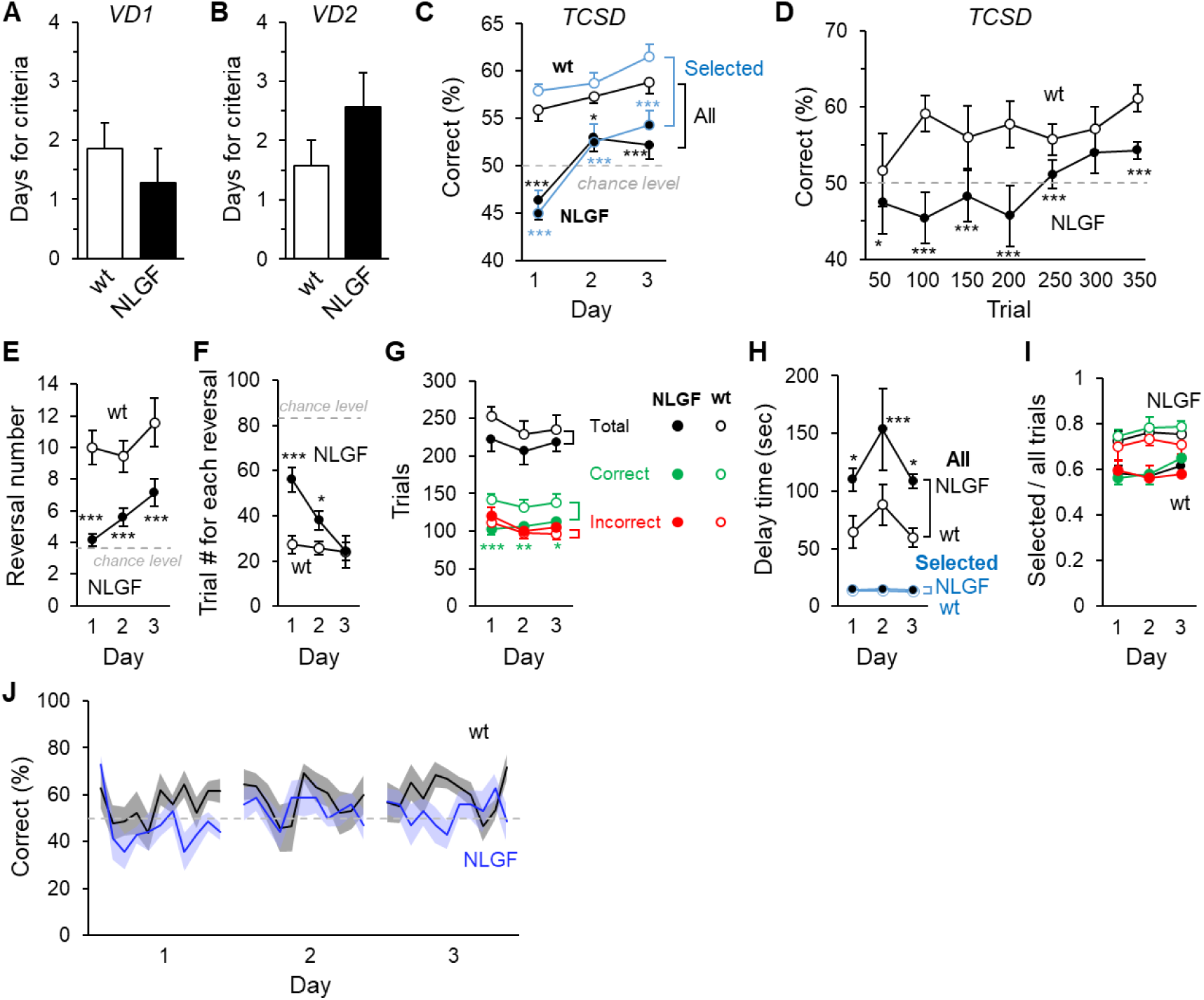
Male *App^NL−G−F^* mice exhibit impaired working memory in the TCSD task at 5 months of age. (**A, B**) The number of days required to reach the performance criterion for the VD1 (**A**) and VD2 (**B**) tasks was comparable between male *App^NL−G−F^* and wild-type (wt) mice. (**C**) TCSD task results. Male *App^NL−G−F^* mice exhibited lower correct rates compared to WT mice. Blue-outlined plots represent selected trials that met the criteria for actual delay times. (**D**) Performance over time. Correct rates for each 50-trial bin during the first 350 trials revealed poorer performance in *App^NL−G−F^* mice compared to WT mice. (**E**) Reversals. The number of reversals per day was reduced in *App^NL−G−F^* mice. (**F**) Trial efficiency. *App^NL−G−F^* mice required more trials per reversal on average compared to WT mice. (**G**) Trial outcomes. *App^NL−G−F^* mice exhibited a lower number of correct trials, while the number of incorrect trials remained comparable between groups. (**H**) Touch intervals. *App^NL−G−F^* mice showed an increased average time between touches. However, this difference was not significant when analyzing selected trials that met the criteria for actual delay times. (**I**) Ratio of selected trials to total trials. The ratio of selected trials to total trials was similar between groups. (**J**) Correct average rates for each 10-trial bin across daily trials. The graph shows results from the first 11 bins of each day. Within a given day, the correct rate remained unchanged. Two-way repeated ANOVA with Bonferroni post-hoc tests. \*\*\**p* < 0.0001; \**p* < 0.01. *n* = 7 per genotype.

In the TCSD task, although the inter-trial interval was set to 5 sec, mice could spend any amount of time between trials. Indeed, the delay time from touching a panel in one trial to touching a panel in the next was significantly longer in *App^NL-G-F^* mice (Fig. 8H; *F*_1,12_ = 8.350, *p =* 0.0136), a result consistent with the findings from the DNMTP task. To exclude the potential influence of longer delay duration, we reanalyzed only the trials with delay times of 20 sec or less. More than half of the trials met this criterion (Fig. 8I), and the delay times in these selected trials were comparable between *App^NL-G-F^* and wild-type mice (Fig. 8H, blue-outlined circles). However, the correct rate remained significantly lower in *App^NL-G-F^* mice for these trials (*F*_1,12_ = 33.381, *p* < 0.0001) (Fig. 8C, blue-outlined circles). Moreover, there was no decline in correct rates across trials within each day, suggesting that motivation remained constant throughout the tasks (paired *t*-test: wild type, *p =* 0.712; *App^NL-G-F^*, *p =* 0.910) (Fig. 8J). Together, these results indicate that the Operant House can detect early working memory deficits in *App^NL-G-F^* mice as early as 5 months old using two independent tasks, DNMTP and TCSD.

### *App^NL-G-F^* mice exhibit abnormal synaptic changes in the mPFC at 5 months of age

Given the impaired working memory detected in 5-month-old *App^NL-G-F^* mice, we next examined whether any morphological or functional changes were present at this stage. Immunohistochemical analyses revealed the accumulation of amyloid β plaques in the mPFC of *App^NL-G-F^* mice (Fig. 9A). In brain regions devoid of amyloid β plaques, the immunoreactivity of presynaptic markers for inhibitory synapses (VGAT: vesicular GABA transporter) was reduced in mPFC sections of *App^NL-G-F^* mice (Fig. 9B, C), while markers for excitatory synapses (VGluT1: vesicular glutamate transporter 1) remained unaffected. Furthermore, the immunoreactivity of both VGluT1 and VGAT was significantly lower inside amyloid β plaques compared to regions outside the plaques (Fig. 9D) in *App^NL-G-F^*mice. These findings suggest that both excitatory and, more prominently, inhibitory synapses are affected in *App^NL-G-F^* mice at 5 months of age.

**Fig. 9.**
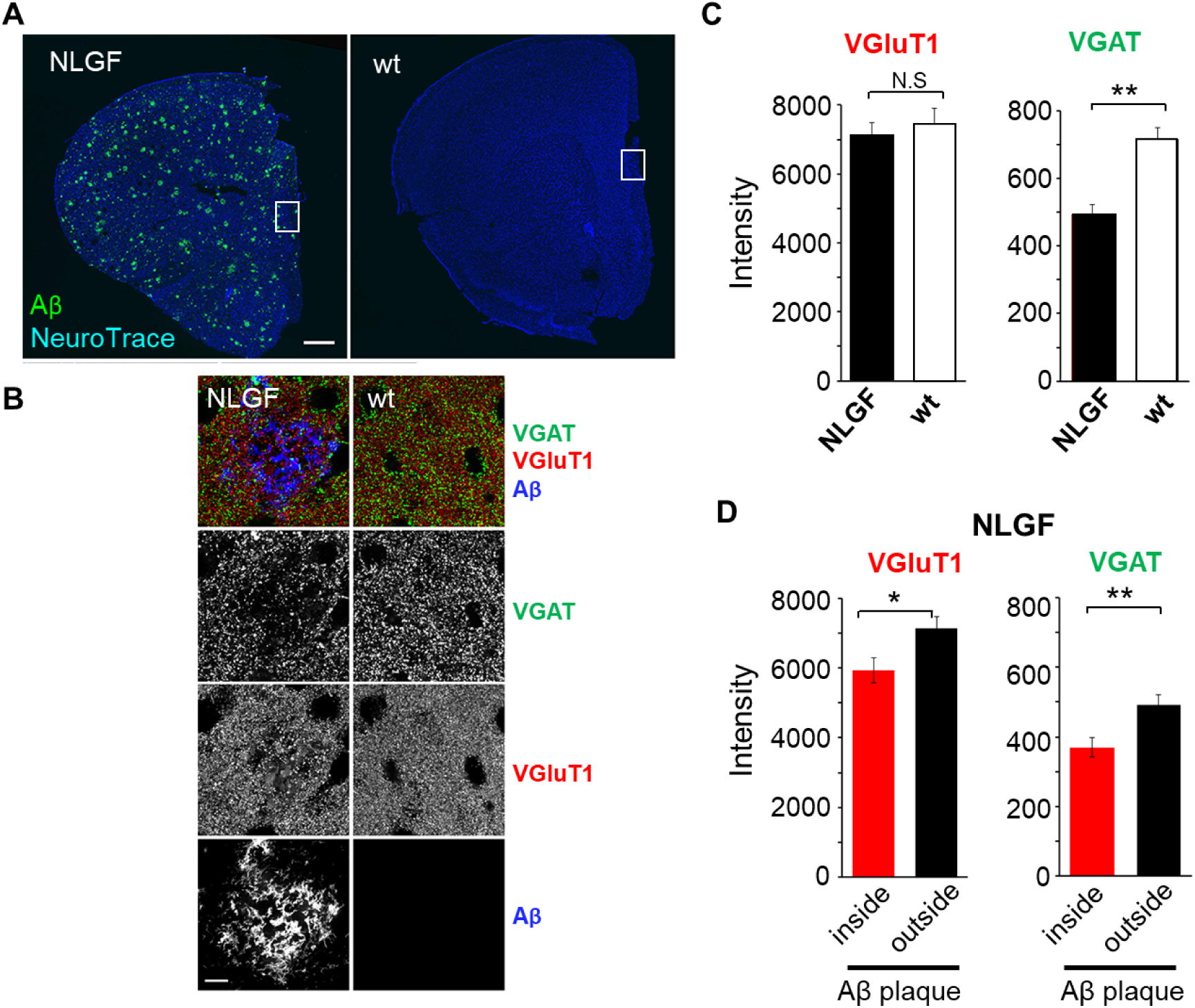
Reduced inhibitory and excitatory presynaptic markers in the mPFC of *App^NL−G−F^* mice at 5 months of age. (**A**) Representative images showing amyloid β plaque accumulation (Aβ) in the mPFC of male 5-month-old *App^NL−G−F^* (NLGF) and their wild-type (wt) littermates. Neurons were stained with Neurotrace. Scale bar, 500 μm (**B**) Enlarged images of the regions indicated by squares in (**A**), immunostained for presynaptic markers VGAT (green), VGluT1 (red), and Aβ (blue). Scale bar, 10 μm. (**C**) Quantification of average fluorescence intensities for VGluT1 (left) and VAT (right) in regions outside of Aβ plaques (defined as clusters of Aβ immunoreactivity > 50 μm^2^). VGAT intensities were decreased in *App^NL−G−F^*group, whereas VGluT1 intensities remained unchanged. (**D**) Average fluorescence intensities for VGluT1 (left) and VGAT (right) were both significantly decreased inside (red) versus outside (black) Aβ plaques in the mPFC of *App^NL−G−F^* mice. Student’s t-test: \**p* < 0.05; \*\**p* < 0.01. *n* = 23-24 regions from *n* = 3 mice for each genotype.

To investigate functional changes in synaptic activity, we performed electrophysiological recordings on acute mPFC slices from 5-month-old *App^NL-G-F^* mice. Basal electrophysiological properties of pyramidal neurons, including input resistance and regular firing rate in response current injection, were comparable between wild-type and *App^NL-G-F^* pyramidal neurons in mPFC slices (Fig. 10A). The amplitudes of miniature excitatory and inhibitory postsynaptic currents (mEPSCs and mIPSCs) were unaffected (Fig. 10B). Similarly, the average ratio of IPSC to EPSC amplitudes (E/I ratio) evoked by local stimulation showed no significant differences in *App^NL-G-F^* pyramidal neurons (Fig. 10C). However, *App^NL-G-F^* pyramidal neurons exhibited greater variability in mIPSC frequency (Fig. 10B, Levene’s test, *p =* 0.0157) and in the E/I ratio (Fig. 10C, Levene’s test, *p <* 0.00001). These results are consistent with the observed preferential reduction in VGAT signals observed in these mice (Fig. 9C). Since neurons surrounding amyloid plaques are selectively impaired in the early stages of Alzheimer’s disease (Busche et al., 2008; Busche et al., 2012), these findings suggest that only a subset of inhibitory neurons is affected, leading to increased variability in inhibitory synaptic functions in the mPFC of 5-months-old *App^NL-G-F^* mice.

**Fig. 10.**
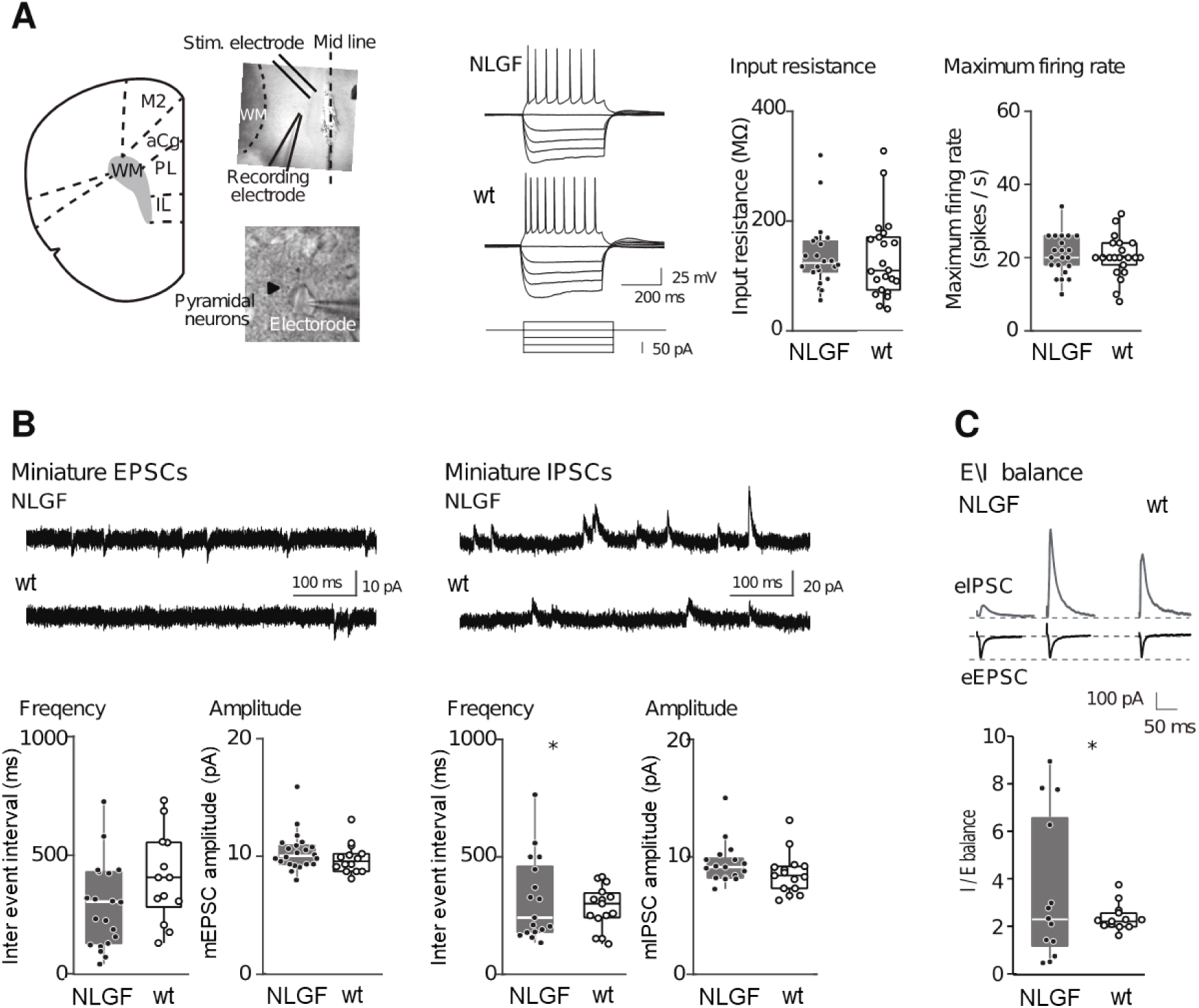
Increased variability in excitatory―inhibitory balance in *App^NL−G−F^* mice at 5 months of age. (**A**) Whole-cell patch-clamp recordings were performed on pyramidal neurons in acute mPFC slices from male 5-month-old *App^NL−G−F^* (NLGF) and wild-type (wt) mice (left). Representative regular firing patterns induced by current injections are shown (middle). Averaged input resistance and maximum firing rates of *App^NL−G−F^* and wild-type pyramidal neurons are presented as mean± SEM (right). (**B**) Frequency and amplitude of miniature EPSCs (mEPSCs) and miniature IPSCs (mIPSCs) in pyramidal neurons. Representative traces are shown (upper). Averaged frequency and amplitude for mEPSC and mIPSC are presented as mean ± SEM. Increased variance in inter-event intervals of mIPSCs was observed in pyramidal neurons from *App^NL−G−F^* mice (Levene’s test, *p =* 0.0157, *n* = 17 cells from *n* = 3 *App^NL−G−F^* mice and 15 cells from *n* = 4 wild-type mice). (**C**) The ratio of evoked IPSC amplitude to EPSC amplitude, with stimulus intensity set to evoke approximately 100 pA for EPSCs, is shown. The distribution for E/I ratio for individual cells is presented as mean ± SEM. Increased variability in excitatory-inhibitory balance was observed in *App^NL−G−F^* mice (Levene’s test; *p* < 0.00001, *n* = 13 cells from *n* = 3 mice for each genotype).

Interestingly, tetanic stimulation (100 pulses at 100 Hz) applied to Schaffer collaterals elicited long-term potentiation (LTP) of excitatory synapses between Schaffer collaterals and CA1 pyramidal neurons in acute dorsal hippocampal slices from both wild-type and *App^NL-G-F^* mice (Supplementary Fig. S7). These findings suggest that LTP is unlikely to contribute to the observed impairments in working memory in *App^NL-G-F^* mice.

## Discussion

In this study, we developed the Operant House, a custom-built, flexible, and programable device designed for automated operant conditioning tests within the home cages of mice. Its compact size (approximately 30 cm on each side) and cost-effectiveness make it suitable for conducting operant conditioning experiments on a large number of animals with minimal experimenter intervention. While several similar home-cage-based operant conditioning devices have been developed recently with comparable goals (Poddar et al., 2013; Pineno, 2014; Devarakonda et al., 2016; Nguyen et al., 2016; Francis and Kanold, 2017; Lee et al., 2020; Matikainen-Ankney et al., 2021), these devices have limitations in the range of operant tasks they can perform. Specifically, they rely on visual stimuli presented at fixed locations and support nose poking at only two or three predetermined positions. In contrast, the Operant House offers an unlimited range of configurations for visual stimuli and nose poke patterns, achieved by simply modifying the program and mask pattern on the LCD monitor. Furthermore, replacing the chamber wall enables an easy transition between nose poke tasks and lever press tasks—a unique feature of the Operant House. Wild-type mice demonstrated rapid learning in a lever-press operant conditioning task within a days, a rate approximately eight times faster than previously reported using commercial devices (Supplementary Fig. S1) (Yu et al., 2009). Additionally, the Operant House employs an open-source Python program that can run on a personal computer (version 1) or a more economical single-board computer, such as Raspberry Pi (version 2). The software is easily customizable by following the official instruction website if user has basic python coding knowledge. Protocols for additional tests, including tilt discrimination, probabilistic reversal learning, pattern separation, and delayed alternation tasks, are continuously updated and freely available on the project website (https://operanthouse.sakura.ne.jp/index.html). The Operant House software supports remote operation, email notifications of daily results, continuous video recording via an IR camera during task execution, and real-time monitoring on computers or smartphones. It can export results in text or CSV formats for post-hoc analysis. Additionally, the software includes a TTL signal output and timestamp functionality, enabling seamless integration with external devices for experiments involving optogenetics. Furthermore, the Operant House is compatible with the UCLA Miniscope when the Raspberry Pi is replaced with a computer connected to the Miniscope. Notably, we successfully conducted in vivo Ca²⁺ recording using the UCLA Miniscope during a visual discrimination task using the Operant House (data not shown). These features make the Operant House a highly versatile, easy-to-construct, and user-friendly tool, well-suited for a broad range of operant conditioning experiments.

As a proof of concept, we evaluated working memory in mice using two tests in the Operant House: the DNMTP and TCSD tests. Spatial working memory in rodents is commonly assessed with the T-maze alternation test (Deacon and Rawlins, 2006). In this test, mice must choose between a previously visited arm and a goal arm, the latter offering a food reward. A simplified version of this task measures spontaneous alternations in open arms without blocking entries (Sarnyai et al., 2000). However, T-maze alternation tests may be confounded by spatial novelty preference, a passive form of short-term memory (Sanderson and Bannerman, 2012). For example, T-maze rewarded alternation performance was unaffected by the loss of NMDA receptors in the prefrontal cortex, an area critical for working memory, whereas DMTP tasks with five visual cues and nose-poking revealed significant impairments (Kilonzo et al., 2021). Similarly, the DNMTP test in the Operant House, which utilizes three positions for visual cues and nose pokes, led to increased c-Fos expression in the prefrontal cortex following task completion. Furthermore, prefrontal cortex lesions impaired performance in this test. These results indicate that the DNMTP test robustly measures prefrontal cortex-dependent working memory in mice.

The *App^NL-G-F^* mouse, a knock-in model that does not overexpress mutant genes, exhibits relatively mild neurological phenotypes, particularly in the early stages. While some studies have reported abnormalities at six months of age in tasks such as the Morris water maze and context fear conditioning (Mehla et al., 2019), most studies have shown normal performance (Saito et al., 2014; Sakakibara et al., 2018; Latif-Hernandez et al., 2019; Saifullah et al., 2020; Kundu et al., 2021). In Alzheimer’s disease, working memory impairment is one of the earliest symptoms during the preclinical stage (Langbaum et al., 2013; Reiman et al., 2016; Weintraub et al., 2018). However, studies of working memory in mouse models of Alzheimer’s disease are limited and controversial. Studies using the Y-maze on *App^NL-G-F^* mice at six months of age have shown mixed results: some report impairments (Saito et al., 2014; Kundu et al., 2021), while others do not (Whyte et al., 2018). Interestingly, studies showing no impairments (Whyte et al., 2018) used male mice, while impairments were observed only in females in another study (Kundu et al., 2021). Although the reasons remain unclear, it is notable that the prevalence of Alzheimer’s disease in humans is approximately twice as high in women as in men. Furthermore, differences in the sex of the experimenter have been suggested as a factor influencing Y-maze results (Whyte et al., 2018). Thus, abnormalities observed only in female *App^NL-G-F^* mice by the DNMTP test in the Operant House, which is conducted without experimenter intervention, are particularly significant. Conversely, impairments in working memory were reported in male *App^NL-G-F^* mice at 4–6 months of age by the location discrimination (LD) test, in which mice choose between two visual stimuli from six locations (Saifullah et al., 2020). Similarly, the TCSD test in the Operant House, which requires mice to choose between two sides based on five visual stimuli, revealed abnormalities in male *App^NL-G-F^* mice at 5 months of age. These findings suggest that the DNMTP and TCSD (LD) tests may assess distinct aspects of working memory or other cognitive functions, such as cognitive inflexibility. The ability to assess working memory in *App^NL-G-F^* mice under natural circadian rhythms (at night) without experimenter intervention highlights the utility of the Operant House, particularly for models exhibiting mild behavioral phenotypes.

Interestingly, while functional synaptic impairments were observed in the mPFC of *App^NL-G-F^* mice at 5 months of age, LTP in the hippocampus, a brain region critical for episodic and spatial memory, remained intact (Supplementary Figure S7). These findings suggest the distinct regional effects of synaptic dysfunction in the early stages of Alzheimer’s disease models. Recent advancements in techniques such as optogenetics and *in vivo* Ca^2+^ imaging have provided valuable insights into the circuit mechanisms underlying working memory (Shirai and Hayashi-Takagi, 2017; Aharoni and Hoogland, 2019). Since Ca^2+^ imaging, optogenetics and operant conditioning can be seamlessly integrated, the Operant House holds promise as a powerful platform to investigate circuit mechanisms underlying working memory deficits in disease models.

## Acknowledgments

We extend our gratitude to Drs. Takashi Saito, Takaomi C. Saido, and Yoichiro Abe for generously providing us with *App^NL-G-F^* mice. This work was supported by the MEXT KAKENHI (Grant Numbers 20H05628 to M.Y. and 22H05158 to A.I-I.) and AMED (Grant Numbers JP24zf0127010 to M.Y and JP19gm6310001 to A.I-I).

## References

Aharoni D, Hoogland TM (2019) Circuit Investigations With Open-Source Miniaturized Microscopes: Past, Present and Future. Front Cell Neurosci 13:141.

Aoki S, Liu AW, Zucca A, Zucca S, Wickens JR (2015) Role of Striatal Cholinergic Interneurons in Set-Shifting in the Rat. J Neurosci 35:9424–9431.

Benner S, Endo T, Kakeyama M, Tohyama C (2015) Environmental insults in early life and submissiveness later in life in mouse models. Front Neurosci 9:91.

Benner S, Endo T, Endo N, Kakeyama M, Tohyama C (2014) Early deprivation induces competitive subordinance in C57BL/6 male mice. Physiol Behav 137:42–52.

Busche MA, Chen X, Henning HA, Reichwald J, Staufenbiel M, Sakmann B, Konnerth A (2012) Critical role of soluble amyloid-β for early hippocampal hyperactivity in a mouse model of Alzheimer’s disease. Proc Natl Acad Sci U S A 109:8740–8745.

Busche MA, Eichhoff G, Adelsberger H, Abramowski D, Wiederhold K, Haass C, Staufenbiel M, Konnerth A, Garaschuk O (2008) Clusters of hyperactive neurons near amyloid plaques in a mouse model of Alzheimer’s disease. Science 321:1686–1689.

Codita A, Gumucio A, Lannfelt L, Gellerfors P, Winblad B, Mohammed AH, Nilsson LN (2010) Impaired behavior of female tg-ArcSwe APP mice in the IntelliCage: A longitudinal study. Behav Brain Res 215:83–94.

Deacon RM, Rawlins JN (2006) T-maze alternation in the rodent. Nat Protoc 1:7–12.

Devarakonda K, Nguyen KP, Kravitz AV (2016) ROBucket: A low cost operant chamber based on the Arduino microcontroller. Behav Res Methods 48:503–509.

Dudchenko PA (2004) An overview of the tasks used to test working memory in rodents. Neurosci Biobehav Rev 28:699–709.

Endo T, Maekawa F, Voikar V, Haijima A, Uemura Y, Zhang Y, Miyazaki W, Suyama S, Shimazaki K, Wolfer DP, Yada T, Tohyama C, Lipp HP, Kakeyama M (2011) Automated test of behavioral flexibility in mice using a behavioral sequencing task in IntelliCage. Behav Brain Res 221:172–181.

Francis NA, Kanold PO (2017) Automated Operant Conditioning in the Mouse Home Cage. Front Neural Circuits 11:10.

Han SD, Nguyen CP, Stricker NH, Nation DA (2017) Correction to: Detectable Neuropsychological Differences in Early Preclinical Alzheimer’s Disease: a Meta-Analysis. Neuropsychol Rev 27:326–327.

Horigane SI, Ozawa Y, Zhang J, Todoroki H, Miao P, Haijima A, Yanagawa Y, Ueda S, Nakamura S, Kakeyama M, Takemoto-Kimura S (2020) A mouse model of Timothy syndrome exhibits altered social competitive dominance and inhibitory neuron development. FEBS Open Bio 10:1436–1446.

Horner AE, Heath CJ, Hvoslef-Eide M, Kent BA, Kim CH, Nilsson SR, Alsio J, Oomen CA, Holmes A, Saksida LM, Bussey TJ (2013) The touchscreen operant platform for testing learning and memory in rats and mice. Nat Protoc 8:1961–1984.

Iman IN, Yusof NAM, Talib UN, Ahmad NAZ, Norazit A, Kumar J, Mehat MZ, Jayabalan N, Muthuraju S, Stefaniuk M, Kaczmarek L, Muzaimi M (2021) The IntelliCage System: A Review of Its Utility as a Novel Behavioral Platform for a Rodent Model of Substance Use Disorder. Front Behav Neurosci 15:683780.

Kilonzo K, van der Veen B, Teutsch J, Schulz S, Kapanaiah SKT, Liss B, Katzel D (2021) Delayed-matching-to-position working memory in mice relies on NMDA-receptors in prefrontal pyramidal cells. Sci Rep 11:8788.

Kirova AM, Bays RB, Lagalwar S (2015) Working memory and executive function decline across normal aging, mild cognitive impairment, and Alzheimer’s disease. Biomed Res Int 2015:748212.

Kiryk A, Janusz A, Zglinicki B, Turkes E, Knapska E, Konopka W, Lipp HP, Kaczmarek L (2020) IntelliCage as a tool for measuring mouse behavior - 20 years perspective. Behav Brain Res 388:112620.

Kundu P, Torres ERS, Stagaman K, Kasschau K, Okhovat M, Holden S, Ward S, Nevonen KA, Davis BA, Saito T, Saido TC, Carbone L, Sharpton TJ, Raber J (2021) Integrated analysis of behavioral, epigenetic, and gut microbiome analyses in App(NL-G-F), App(NL-F), and wild type mice. Sci Rep 11:4678.

Kwak C, Lim CS, Kaang BK (2015) Development of a touch-screen-based paradigm for assessing working memory in the mouse. Exp Neurobiol 24:84–89.

Langbaum JB, Fleisher AS, Chen K, Ayutyanont N, Lopera F, Quiroz YT, Caselli RJ, Tariot PN, Reiman EM (2013) Ushering in the study and treatment of preclinical Alzheimer disease. Nat Rev Neurol 9:371–381.

Latif-Hernandez A, Shah D, Craessaerts K, Saido T, Saito T, De Strooper B, Van der Linden A, D’Hooge R (2019) Subtle behavioral changes and increased prefrontal-hippocampal network synchronicity in APP(NL-G-F) mice before prominent plaque deposition. Behav Brain Res 364:431–441.

Lee JH, Capan S, Lacefield C, Shea YM, Nautiyal KM (2020) DIY-NAMIC Behavior: A High-Throughput Method to Measure Complex Phenotypes in the Homecage. eNeuro 7.

Matikainen-Ankney BA et al. (2021) An open-source device for measuring food intake and operant behavior in rodent home-cages. Elife 10.

Mehla J, Lacoursiere SG, Lapointe V, McNaughton BL, Sutherland RJ, McDonald RJ, Mohajerani MH (2019) Age-dependent behavioral and biochemical characterization of single APP knock-in mouse (APP(NL-G-F/NL-G-F)) model of Alzheimer’s disease. Neurobiol Aging 75:25–37.

Nguyen KP, O’Neal TJ, Bolonduro OA, White E, Kravitz AV (2016) Feeding Experimentation Device (FED): A flexible open-source device for measuring feeding behavior. J Neurosci Methods 267:108–114.

Pineno O (2014) ArduiPod Box: a low-cost and open-source Skinner box using an iPod Touch and an Arduino microcontroller. Behav Res Methods 46:196–205.

Poddar R, Kawai R, Olveczky BP (2013) A fully automated high-throughput training system for rodents. PLoS One 8:e83171.

Reiman EM, Langbaum JB, Tariot PN, Lopera F, Bateman RJ, Morris JC, Sperling RA, Aisen PS, Roses AD, Welsh-Bohmer KA, Carrillo MC, Weninger S (2016) CAP--advancing the evaluation of preclinical Alzheimer disease treatments. Nat Rev Neurol 12:56–61.

Remmelink E, Chau U, Smit AB, Verhage M, Loos M (2017) A one-week 5-choice serial reaction time task to measure impulsivity and attention in adult and adolescent mice. Sci Rep 7:42519.

Rossi MA, Yin HH (2012) Methods for studying habitual behavior in mice. Curr Protoc Neurosci Chapter 8:Unit 8 29.

Saifullah MAB, Komine O, Dong Y, Fukumoto K, Sobue A, Endo F, Saito T, Saido TC, Yamanaka K, Mizoguchi H (2020) Touchscreen-based location discrimination and paired associate learning tasks detect cognitive impairment at an early stage in an App knock-in mouse model of Alzheimer’s disease. Mol Brain 13:147.

Saito T, Matsuba Y, Mihira N, Takano J, Nilsson P, Itohara S, Iwata N, Saido TC (2014) Single App knock-in mouse models of Alzheimer’s disease. Nat Neurosci 17:661–663.

Sakakibara Y, Sekiya M, Saito T, Saido TC, Iijima KM (2018) Cognitive and emotional alterations in App knock-in mouse models of Abeta amyloidosis. BMC Neurosci 19:46.

Sanderson DJ, Bannerman DM (2012) The role of habituation in hippocampus-dependent spatial working memory tasks: evidence from GluA1 AMPA receptor subunit knockout mice. Hippocampus 22:981–994.

Sarnyai Z, Sibille EL, Pavlides C, Fenster RJ, McEwen BS, Toth M (2000) Impaired hippocampal-dependent learning and functional abnormalities in the hippocampus in mice lacking serotonin 1A receptors. PNAS 97:14731–14736.

Shirai F, Hayashi-Takagi A (2017) Optogenetics: Applications in psychiatric research. Psychiatry Clin Neurosci 71:363–372.

Ujita W, Kohyama-Koganeya A, Endo N, Saito T, Oyama H (2018) Mice lacking a functional NMDA receptor exhibit social subordination in a group-housed environment. FEBS J 285:188–196.

van Dijk RM, Wiget F, Wolfer DP, Slomianka L, Amrein I (2019) Consistent within-group covariance of septal and temporal hippocampal neurogenesis with behavioral phenotypes for exploration and memory retention across wild and laboratory small rodents. Behav Brain Res 372:112034.

Voikar V, Krackow S, Lipp HP, Rau A, Colacicco G, Wolfer DP (2018) Automated dissection of permanent effects of hippocampal or prefrontal lesions on performance at spatial, working memory and circadian timing tasks of C57BL/6 mice in IntelliCage. Behav Brain Res 352:8–22.

Weintraub S, Carrillo MC, Farias ST, Goldberg TE, Hendrix JA, Jaeger J, Knopman DS, Langbaum JB, Park DC, Ropacki MT, Sikkes SAM, Welsh-Bohmer KA, Bain LJ, Brashear R, Budur K, Graf A, Martenyi F, Storck MS, Randolph C (2018) Measuring cognition and function in the preclinical stage of Alzheimer’s disease. Alzheimers Dement (N Y) 4:64–75.

Whyte LS, Hemsley KM, Lau AA, Hassiotis S, Saito T, Saido TC, Hopwood JJ, Sargeant TJ (2018) Reduction in open field activity in the absence of memory deficits in the App(NL-G-F) knock-in mouse model of Alzheimer’s disease. Behav Brain Res 336:177–181.

Yu C, Gupta J, Chen JF, Yin HH (2009) Genetic deletion of A2A adenosine receptors in the striatum selectively impairs habit formation. J Neurosci 29:15100–15103.

