## Supplemental Figures for "Operant House: A Versatile Open-Source Platform for Automated Operant Conditioning Testing of Mice in Home Cages"

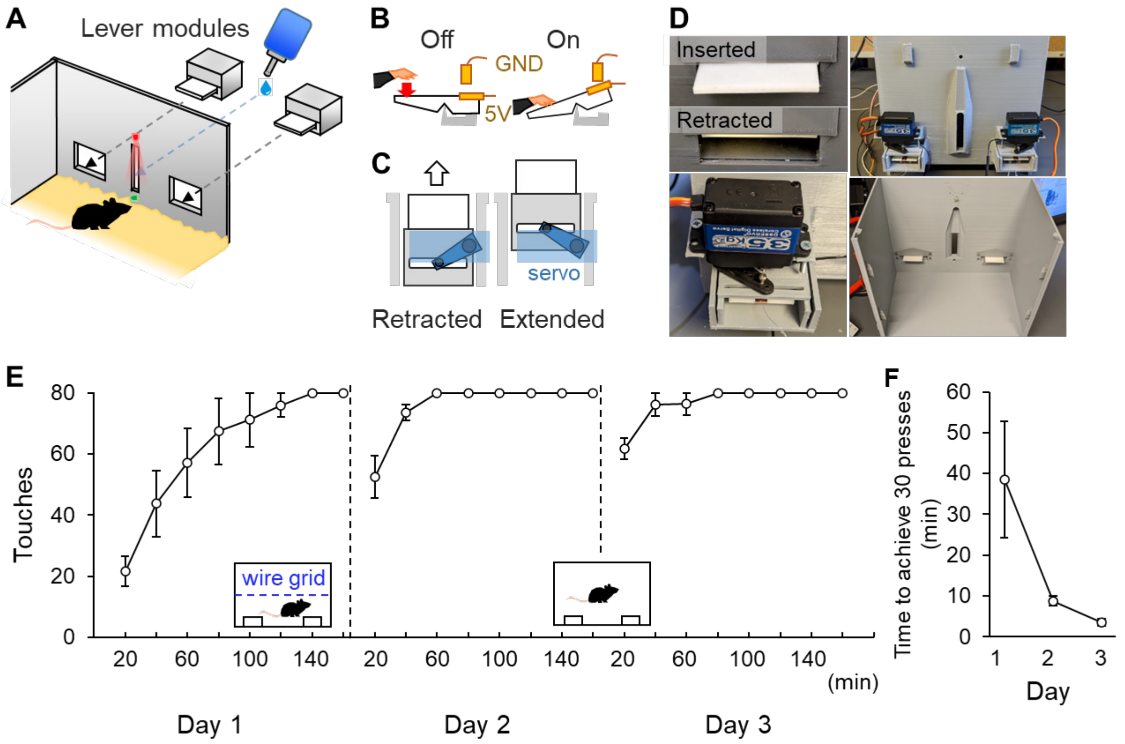


**Supplementary Fig. S1. Related to Fig. 3. Retractable lever modules.**

(**A-C**) Design and schematic representation of the retractable lever module. Two or more lever modules can be installed to the default chamber (**A**). The lever operates using a seesaw-like mechanism and returns to its original position via gravity (**B**), requiring minimum force to push (< 75 mg). A servo motor enables the lever modules slide back and forth (**C**).

(**D**) Pictures of the lever modules. Close-up of images of the inserted and retracted lever (upper-left), an external view of lever modules (upper right), a detailed view of the module's exterior (bottom left), and a view of the levers from inside the chamber (bottom right).

(**E-F**) Lever press performance in wild-type mice (*n* = 5). Number of lever pressed in each time bin (**E**) and elapsed time to achieve 30 presses (**F**).

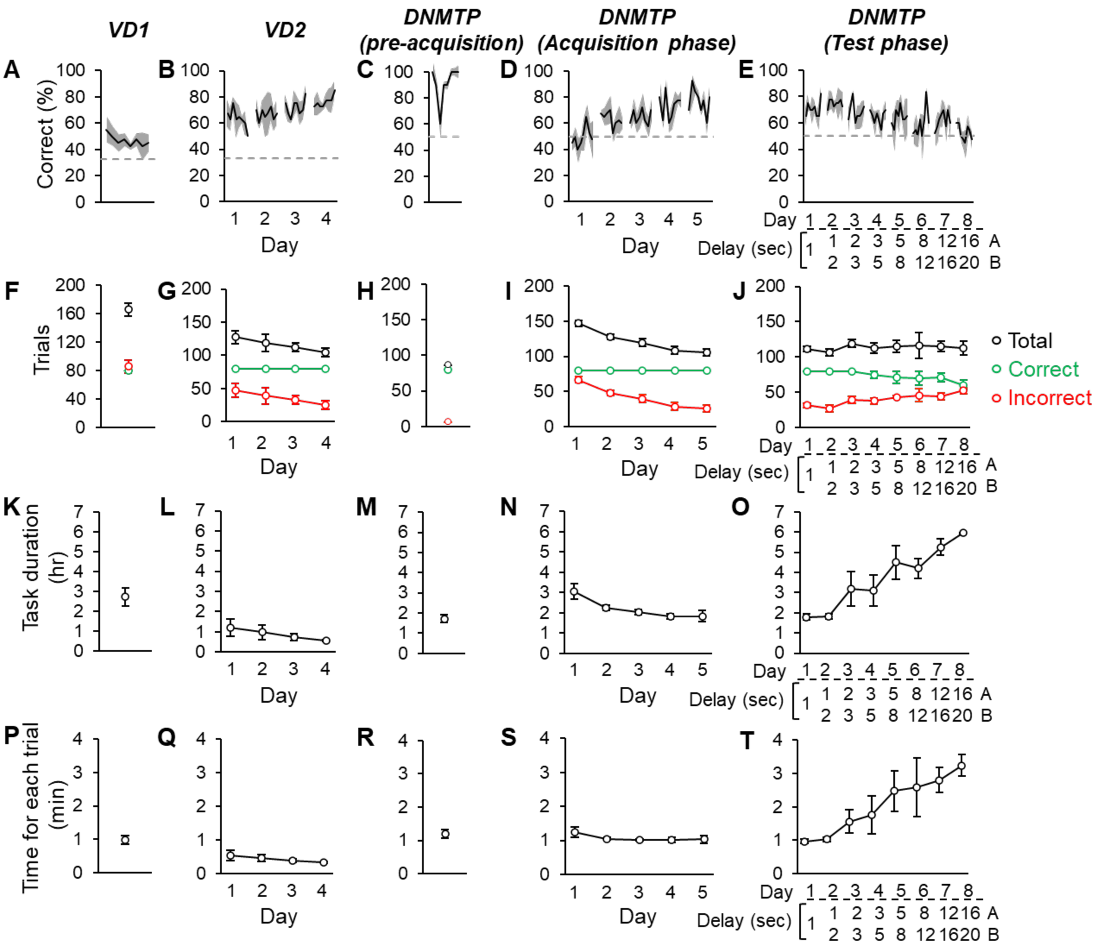


**Supplementary Fig. S2, related to Fig. 5. Additional results from the DNMTP test in wild-type mice.**

(**A-E**) Correct rates for each 10-trial bin across tasks. Graphs display results from the first 8 bins of each day. The average correct rate within each task/day remained relatively constant.

(**F-J**) Total (black circles), correct (green), and incorrect (red) trial numbers for each day.

(**K-O**) Task durations for each day.

(**P-T**) Average time elapsed per trial.

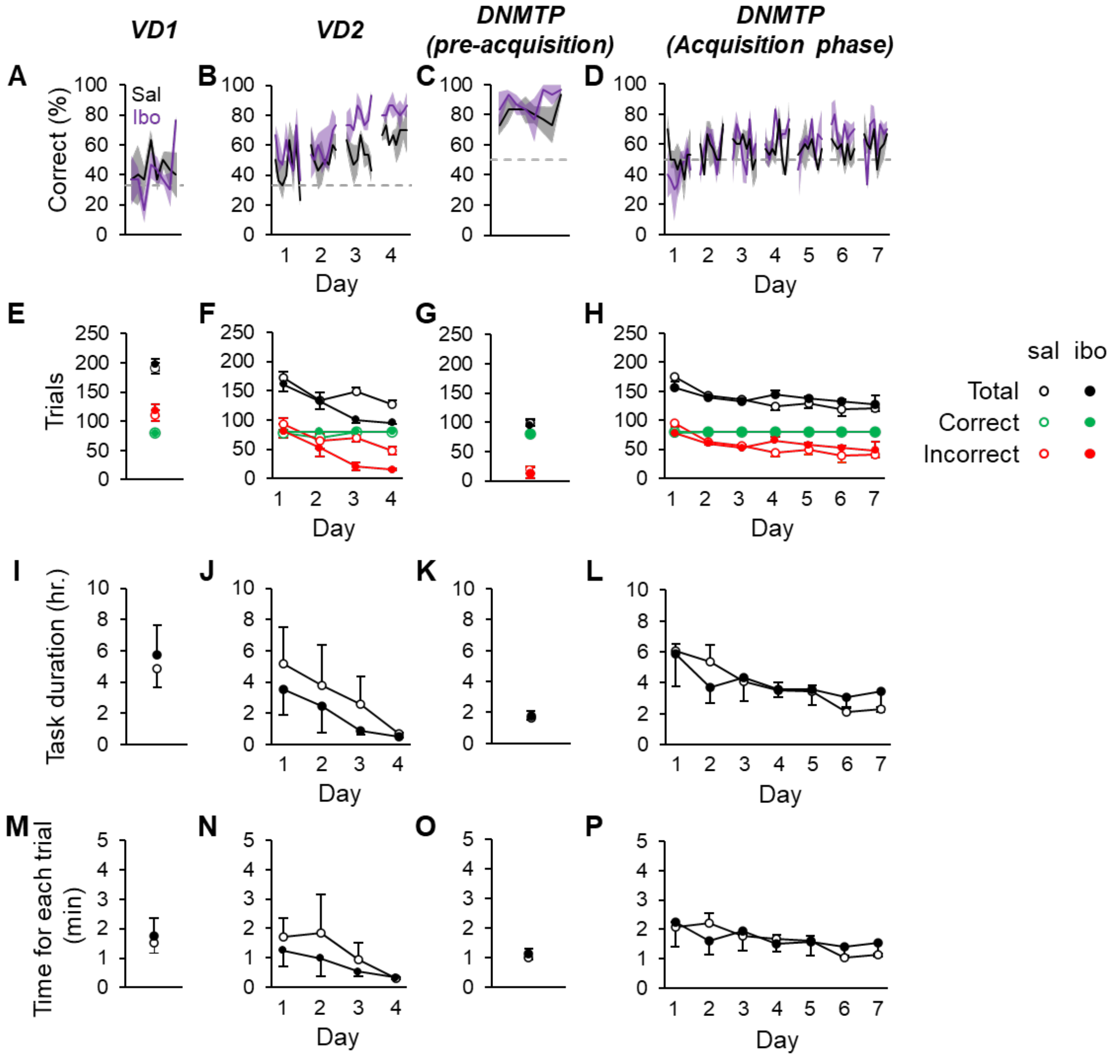


**Supplementary Fig. S3, related to Fig. 5. Additional results of the DNMTP task in wild-type mice with prefrontal cortex lesions induced by ibotenic acid.**

(**A-D**) Correct rates for each 10-trial bin across tasks for saline- and ibotenic acid-treated mice. Graphs show results from the first 8 bins of each day.

(**E-H**) Total (black circles), correct (green), and incorrect (red) trial counts for each day.

(**I-L**) Task duration for each day.

(**M-P**) Average time elapsed per trial.

Statistical analysis was performed using two-way repeated-measures ANOVA with Bonferroni post-hoc tests: ***$p < 0.0001$; **$p < 0.01$; *$p < 0.05$. $n = 3$ mice per group.

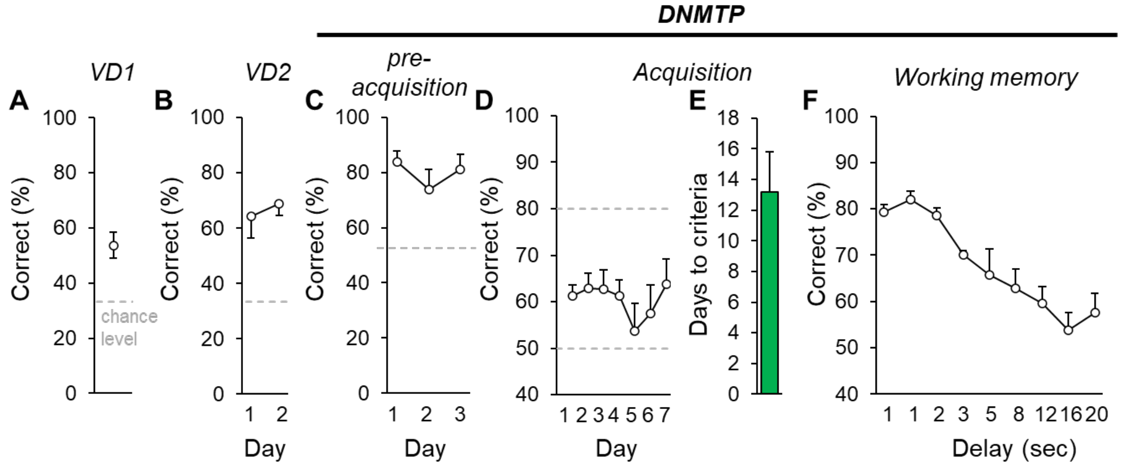


**Supplementary Fig. S4. Results of the DNMTP test using Operant House Version 2.**

(**A-B**) Visual discrimination task results.

(**C**) Pre-acquisition phase of the DNMTP test.

(**D-F**) Acquisition and working memory task results.

Data were obtained from wild-type mice (*n* = 5) in Operant House Version 2.

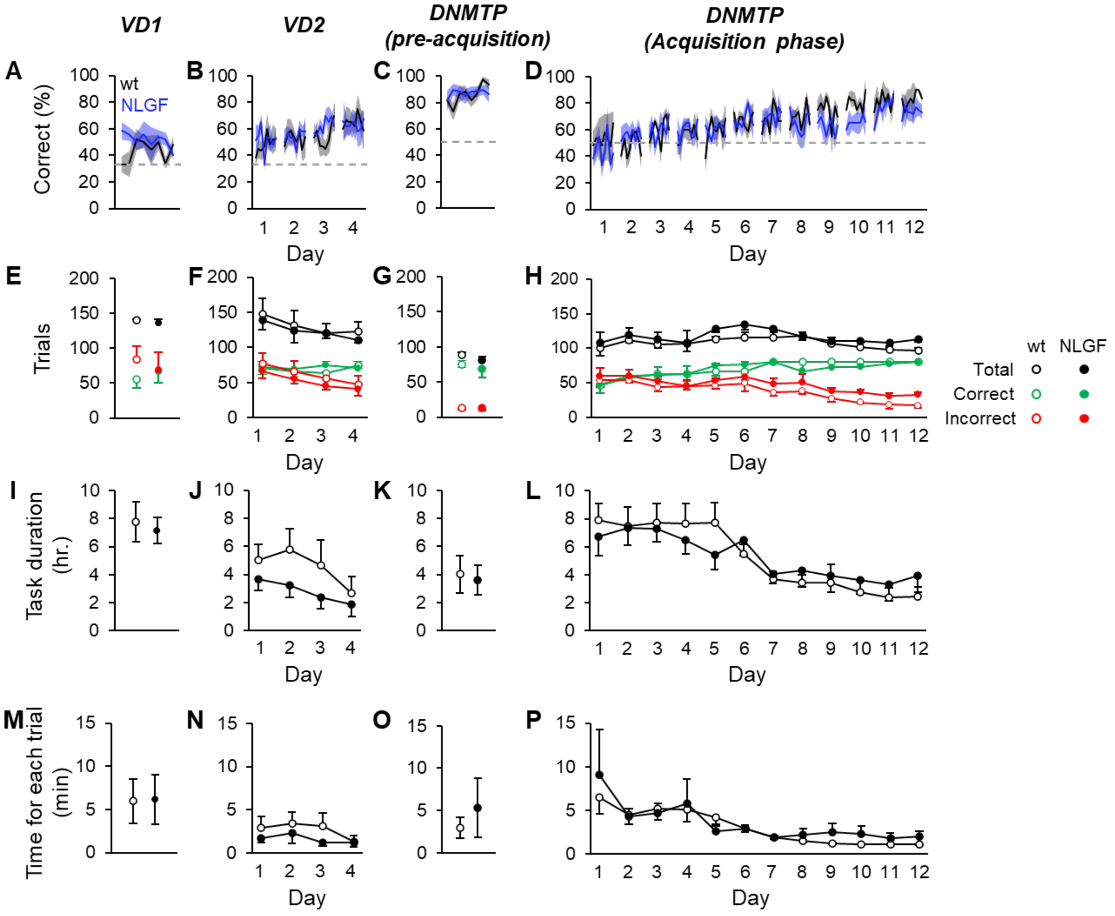


**Supplementary Fig. S5, related to Fig. 8. Additional results of the DNMTP test in female** ***App***$^{NL\text{-}G\text{-}F}$ **mice.**

(**A-D**) Correct rates for each 10-trial bin across different tasks. Results are shown for the first 8 bins recorded each day.

(**E-H**) The total number of trials (black), correct trials (green), and incorrect trials (red) performed each day.

(**I-L**) The trial duration for each day.

(**M-P**) The average time elapsed per trial.

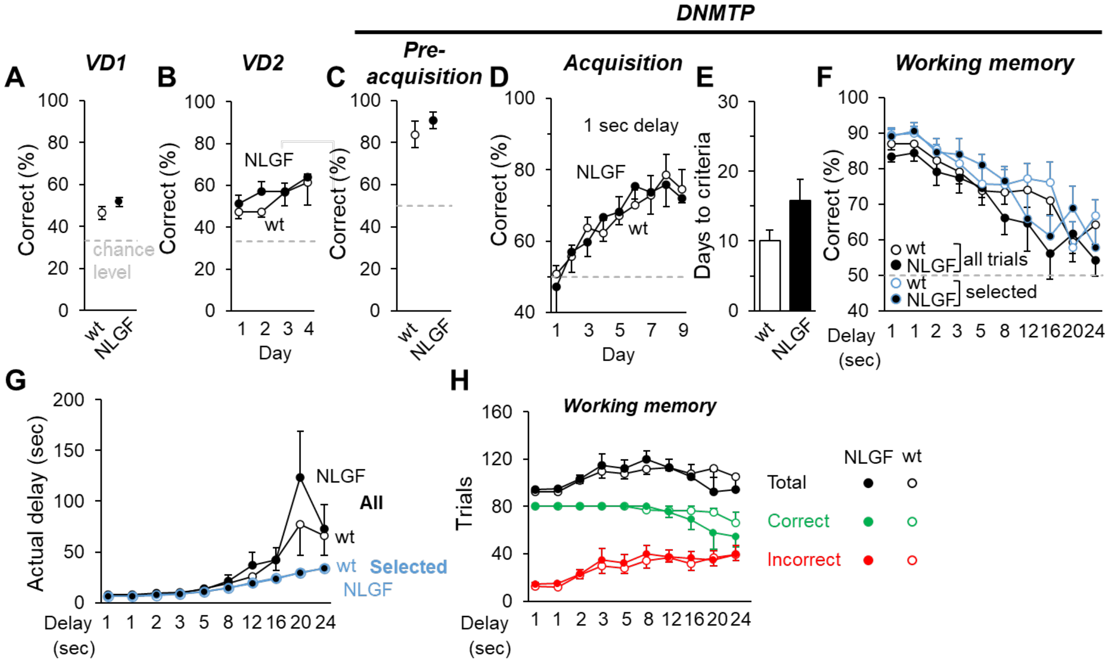


**Supplementary Fig. S6, related to Fig. 6. Male** ***App***$^{NL-G-F}$ **mice exhibited normal working memory in the DNMTP test at 5 months of age.**

(**A**, **B**) Results of VD1 and VD2 tasks in male *App*$^{NL-G-F}$ homozygous mice at 5 months.

(**C-F**) Result of the DNMTP test, including pre-acquisition (**C**), acquisition (**D-E**) and working memory (**F**) phases. No significant differences were observed between genotypes.

(**G**) Actual delay times in trials meeting the criteria for delay were compatible between wild-type and *App*$^{NL-G-F}$ mice.

(**H**) Total numbers of each trial type across all trials. $n$ = 6 for controls and $n$ = 5 for homozygous mice.

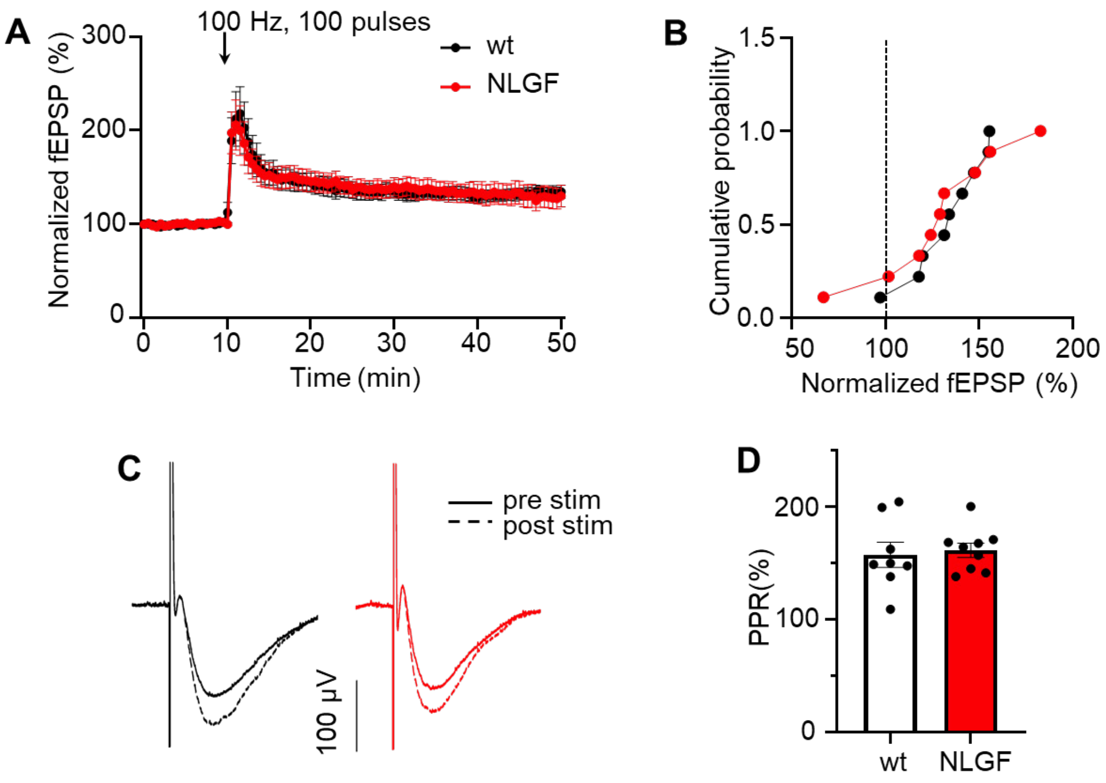


**Supplementary Fig. S7. Related to Fig. 10. No defect in hippocampal long-term potentiation (LTP) in 5-month-old *App*$^{NL-G-F}$ mice.**

(**A**) Representative LTP traces after tetanic stimulation (100 pulses at 100 Hz) of Schaffer collaterals in CA1 pyramidal neurons.

(**B**) Cumulative plots of normalized field excitatory postsynaptic potential (fEPSP) amplitudes. Amplitudes of fEPSPs during the post-stimulation period (40–50 min) were normalized to pre-stimulation values (0–10 min).

(**C**) Representative traces of fEPSPs recorded before and after LTP induction.

(**D**) No differences in paired-pulse ratios (PPR) of fEPSPs between wild-type and *App*$^{NL-G-F}$ mice after LTP induction. PPR was calculated as the ratio of the amplitude of the second pulse-evoked sper genotype.
